# Data selection choices influence the inferred movement patterns of Plasmodium sporozoites in skin

**DOI:** 10.64898/2026.06.29.735005

**Authors:** Sayantan Biswas, Estefania Hurtado, Vitaly V. Ganusov

## Abstract

Motility of *Plasmodium* sporozoites (**SPZs**) in the skin is a key determinant of successful host infection. Earlier studies have described rapid movement of both murine and human SPZs in skin following syringe inoculation. It is typical to classify SPZ trajectories into “motile” and “immotile” and restrict the analysis of movement patterns to motile SPZs. Because criteria to define motile SPZs are dependent on the study and are often qualitative, it remains unclear if sub-selection of motile tracks introduces biases in characterization of SPZ movement in vivo. We processed imaging data (22 movies) from a recent study of movement of *P. falciparum* (**Pf**) and *P. yoelii* (**Py**) SPZ in skin. We proposed a novel metric – maximal spatial spread (**MSS** or *S*) — that is the maximum Euclidean distance between any two recorded positions in a trajectory. We used MSS to classify SPZ trajectories as immotile (*S* ≤ *S*_threshold_) or motile (*S* > *S*_threshold_) for a given threshold value *S*_threshold_. Larger *S*_threshold_ values naturally resulted in a smaller fraction of tracks classified as motile, and subsequently, in an increased overall displacement, instantaneous and mean speeds, decreased mean turning angle, and higher initial slopes of the mean squared displacement (**MSD**) curves. We found that at intermediate values of *S*_threshold_ Pf SPZs had a lower average speed than Py SPZs suggesting that host environment may impact SPZ movement. Both species exhibited a small but statistically significant decline in average speed with time after inoculation but this was also dependent on the *S*_threshold_ value. Our analysis of MSD curves and turning angle distributions suggests that both Pf and Py SPZs undergo correlated random walks – a type of Brownian walk with short-term superdiffusive displacement. By using a novel methodology of hidden Markov models (moveHMM package in R) we found that SPZ movement is best described by three movement states; however, none of these states corresponded to previously described circling gliding. Taking together, our results suggest that inference of SPZ movement patterns depends on the criteria used to define tracks as motile or immotile. Standardized preprocessing criteria are therefore important when comparing motility across *Plasmodium* species, experimental time points, or laboratories. Analysis of turning angle distributions and application of hidden Markov models provided additional metrics to quantify distinct modes of SPZ movement in vivo.

## Introduction

Malaria remains a major global health challenge and continues to cause substantial morbidity and mortality, particularly in regions with high transmission intensity^1–3^. Skin-stage infection begins when Plasmodium sporozoites (**SPZs**) are deposited into the skin during the probing for blood meal by an infected Anopheles mosquito. After inoculation, sporozoites migrates through the dermal tissue, enters blood vessels, and reaches the liver where they invade hepatocytes and form liver stages^1,4^. Because only a fraction of skin-inoculated parasites successfully reach the liver, skin phase of the infection represents a critical bottleneck in malaria transmission^4,5^. SPZs move in skin using substrate-dependent gliding motility driven by an actin–myosin motor complex and associated adhesion proteins such as the thrombospondin-related anonymous protein (**TRAP**)^6,7^. This motility enables parasites to traverse dermal tissue and locate vascular entry points required for dissemination to the liver^8^. Quantitative understanding of SPZ motility in the skin can therefore inform about efficiency at which the parasites cause blood stage infection.

Accurate quantification of SPZ motility is important when evaluating interventions targeting the pre-erythrocytic stages of malaria. Several vaccine and antibody-based strategies aim to prevent infection by interfering with parasite migration before sporozoites reach the liver. Antibodies (**Abs**) directed against the circumsporozoite protein (**CSP**), the dominant SPZ surface protein and the antigen used in the RTS,S and R21 malaria vaccines, have been shown to SPZ motility and prevent parasites from efficiently migrating through the dermis and reaching blood vessels^5,9–11^. In addition, genetic disruption of key components of the parasite gliding technique, including TRAP and associated motor proteins, leads to severe defects in sporozoite motility and dramatically reduces parasite infectivity^6,7^. Consequently, accurate and consistent quantification of sporozoite motility is essential for interpreting parasite migration dynamics and for reliably assessing how genetic or immunological interventions affect parasite movement.

Individual skin-injected SPZs vary dramatically in the type of movement they exhibit; analysis of these movement types often requires to define which parasites are motile and which are immotile. Immotile parasites may be dead and/or dying and are not likely to contribute to blood-stage infection but may be responsible for local skin inflammation; quantifying a fraction of SPZs that are immotile may be useful in evaluating efficacy of SPZ-specific Abs. However, the criteria used to define which SPZs are motile vary substantially between different studies and include a type of gliding behavior, displacement, velocity, or trajectory length (**Table 1**). For example, some in vitro studies defined motile SPZs as those undergoing continuous circular or directed gliding^7,12^; however, such studies did not clearly define quantitative criteria that distinguish motile from immotile SPZs. More recently studies did define thresholds for motile SPZs; for example, Hopp *et al*. ^13^ defined motile SPZs those that had the total displacements more than 2 *µ*m for the whole observation period, and De Korne *et al*. ^14^ defined motile SPZs those with overall displacement of more than 21 pixels (**Table 1**). As far as we are aware, there have been no studies that rigorously investigated the impact of choosing particular metric to define SPZs as motile (or immotile) on inferred movement characteristics. Thresholds to categorize parasites as motile that are too permissive may classify noise or minor positional fluctuations as active movement, whereas overly restrictive thresholds may exclude slowly migrating parasites that may be biologically relevant. Consequently, threshold choice may influence estimates of the fraction of motile parasites and statistical properties of parasite movement programs.

**Table 1:** Example of studies on SPZ motility and the criteria used to define motile SPZs. We reviewed studies of SPZ motility in vitro and in vivo and extracted criteria used in the studies to define which SPZs are deemed as motile. Top studies did not list a specific numerical threshold that defines motile SPZs (denoted by *\**), while bottom studies have listed specific thresholds for motile SPZs. Notations are Pf – P. falciparum, Py – P. yoelii, Pb – P. berghei.

| Study | Parasite | Host system | Motility classification criterion |
| --- | --- | --- | --- |
| Sultan *et al.*<sup>6</sup> | *Pb* | Glass slides | Parasites exhibiting continuous circular gliding over the observation period were classified as motile *. |
| Frischknecht & Matuschewski<sup>7</sup> | *Pb* | Glass/coated substrates | Parasites displaying sustained circular or helical gliding trajectories over time were classified as motile *. |
| Vanderberg & Frevert<sup>5</sup> | *Pb* | Mouse dermis | Parasites showing measurable displacement across sequential imaging frames were classified as motile *. |
| Amino *et al.*<sup>4</sup> | *Pb* | Mouse dermis | Parasites maintaining continuous migration trajectories during intravital imaging were classified as motile *. |
| Münter *et al.*<sup>12</sup> | *Pb* | Glass surface | Parasites that were gliding and/or circling were defined as motile *. |
| Govindasamy *et al.*<sup>15</sup> | *Pb* | Glass surface | Motility was quantified by the percentage of sporozoites that glide and the number of circles they make; the paper reports averages such as $15.7 \pm 1.7$ circles/sporozoite in 180 s in one condition. |
| Hopp *et al.*<sup>13</sup> | *Pb* | Mouse dermis | Trajectories with total displacement $\geq 2 \mu\text{m}$ over the observation period were classified as motile. |
| Hopp *et al.*<sup>16</sup> | *Pb* | Mouse dermis | Trajectories with total displacement $\geq 2 \mu\text{m}$ over the observation period were classified as motile. |
| Hopp *et al.*<sup>17</sup> | *Pf*, *Pb*, *Py* | Mouse dermis and human skin graft xenograft | Trajectories with total displacement $\geq 2 \mu\text{m}$ over the observation period were classified as motile. |
| Klug *et al.*<sup>18</sup> | *Pb* | Glass surface | Parasites completing $\geq 1$ circular trajectory within 5 minutes were classified as motile |
| De Korne *et al.*<sup>14</sup> | *Pb* | Mouse dermis | Trajectories with displacement exceeding a threshold ($\sim 21$ pixels) over the observation period were classified as motile. |
| Liu *et al.*<sup>19</sup> | *Pb* | 3D hydrogel matrix | Motile states defined by instantaneous velocity $\geq 15 \mu\text{m/s}$; velocities below this threshold were classified as pauses or non-motile states. |

In this study, we propose a novel metric to define motile SPZs – the maximal spatial spread that is the maximal Euclidian displacement across all positions of a given SPZ during imaging. We processed multiple movies from intravital imaging experiments of Pf and Py SPZs moving in murine skins at different times after syringe inoculation^13,17^; for total 22 datasets we found that maximal spatial spread varies up to 1,000 fold between individual SPZs for the two species. Using different threshold values of the maximal spatial spread to define motile/immotile SPZs we found that higher thresholds led to higher average SPZ speed, smaller average TAs, and more directed displacement characterized by a faster increase in mean squared displacement with time. However, this came at the cost of a smaller number of trajectories defined as motile. Interestingly, higher threshold values resulted in smaller decline in average SPZ speed with time since inoculation illustrating that choice of which SPZs are motile may influence data interpretation. We suggest that analysis of distribution of maximal spatial spread of individual SPZs may be used to define threshold values to classify SPZs as motile or immotile.

## Materials and methods

### Intravital imaging of Plasmodium sporozoites in the skin

Previous studies used fluorescently labeled Plasmodium sporozoites (**SPZs**) to track their movement in murine skin^13,17^. Specifically, Hopp *et al*. ^17^ injected the 0.2 *µ*L of SPZ suspension (SPZs were isolated from salivary glands of infected mosquitoes) intradermally into the mouse ear pinna (**Figure 1**A). The authors then imaged the exposed skins using intravital microscope with 3-5 z-slices of the total depth of 30 − 50 *µ*m at frequency 1 imaging volume per sec. Imaging was done at different times after SPZ inoculation (e.g., 5, 10, 20, 30, 60, and 120 min) each for a duration of 4 min (or about 240 time frames). The data were exported as TIFF files using maximum projection on z-plane as one z-slice of about 2 *µ*m thick.

**Figure 1:**
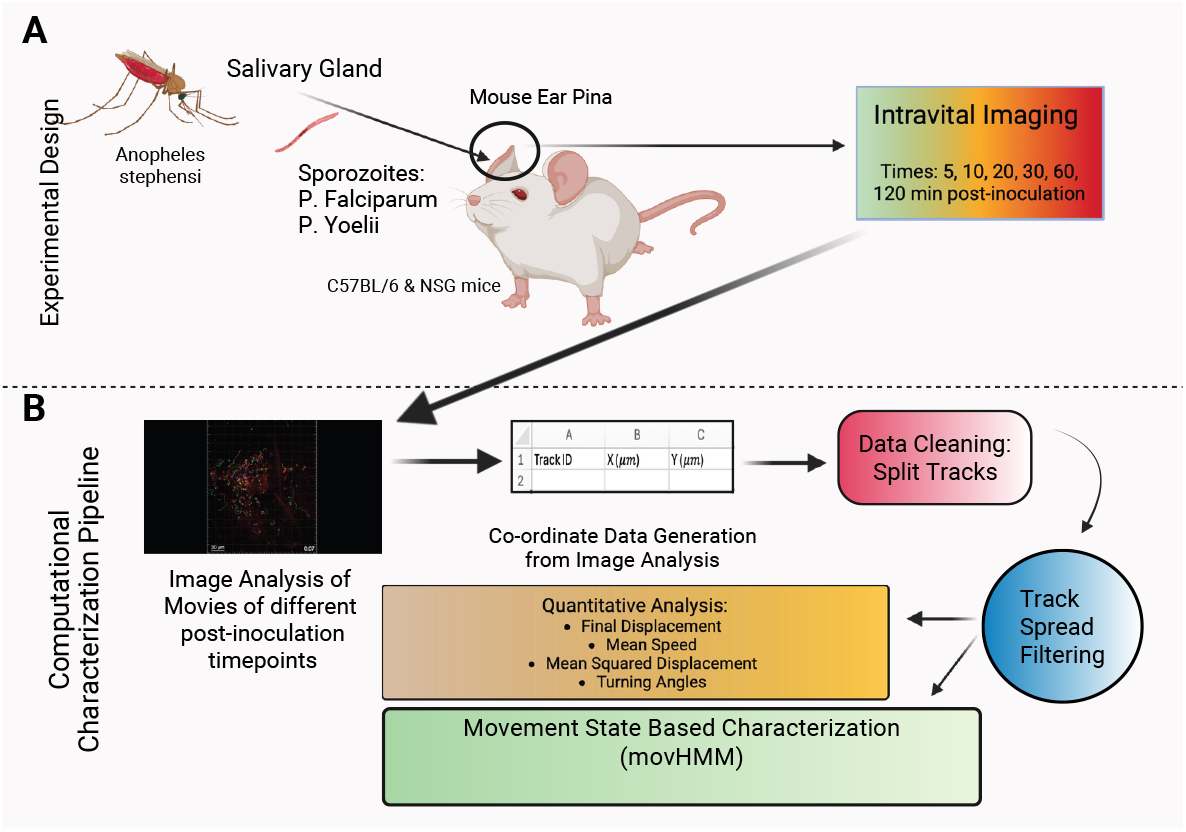
Experimental design and computational analysis workflow for quantifying *Plasmodium* sporozoite motility in murine skin. **A**: Sporozoites of *Plasmodium falciparum* and *P. yoelii* were isolated from the salivary glands of *Anopheles stephensi* mosquitoes and injected into the ear pinnae of Swiss Webster mice ^13^. Intravital fluorescence imaging was performed at multiple time points post-inoculation (5, 10, 20, 30, 60, and 120 min). **B**: the computational characterization pipeline for quantitative analysis of final displacement, mean and instantaneous speeds, mean squared displacement (**MSD**), and turning angles, and state-based characterization using hidden Markov models^20,21^. Coordinate data generated from Imaris-based analysis of movies (**Movies 1–2**) were cleaned by splitting tracks with missing time frames, followed by spatial-spread filtering to classify trajectories as motile or immotile SPZs. All analyses were done with motile trajectories.

### Quantifying SPZ positions over time from movies

#### Analysis of movies

Hopp *et al*. ^17^ performed multiple experiments tracking movement of different species of Plasmodium SPZs. While SPZ coordinate data were not available in the original paper, Dr. Christina Hopp shared with us (via upload to a google drive folder) TIFF-based movies from 7 different experiments: 3 experiments with P. falciparum (experiments labeled as 150724, 150727, and 160323) and 4 with P. yoelii (experiments labeled as 140222, 140223, 150728, 150723); each experiment had the movies for different times after SPZ inoculation. In total, we received 126 TIFF files: 48 were on movement of *P. falciparum* (**Pf**) and 78 were on movement of *P. yoelii* (**Py**). Upon initial review, we found 20 of movies for Py to be corrupted (probably due to poor transfer to Google Drive), and a further 6 Pf and 3 Py movies to contain only blank frames without any visible sporozoites. Excluding those 29 unusable files left us with total of 97 trackable movies.

Previously, Hopp *et al*. ^17^ used software ICY (https://icy.bioimageanalysis.org/) to quantify SPZ positions in the movies. Here to track SPZ positions we opted for using Imaris (https://imaris.oxinst.com/), a proprietary software that allows automatic tracking of objects and manual correction of the identified tracks. We found, however, that automatic tracing of SPZ positions in Imaris was not very efficient and required careful manual editing of the tracks; this precluded us from the analysis of all available movies. We initially analyzed 48 movies: 20 for Pf and 28 for Py. Upon further inspection, out of these 48 movies we restricted further detailed analysis to a subset of 22 movies where we were most confident about identifying all SPZ tracks accurately: 11 movies for Pf SPZs (from experiments 150724, 150727, 160323) and 11 for Py SPZs (from experiments 140222, 140223, 150723). From each experiment we rigorously analyzed movies for different times after SPZ inoculation to ensure equal representation of each species across all experimental time points and to keep the overall tracking workload manageable.

To identify and track SPZs, we developed a custom processing pipeline in Imaris (**Figure 1**B). In order to preserve original intensities we performed no image pre-processing, such as thresholding or filtering. The tracking of SPZs was first automatically performed by the software, which was then followed by the manual editing of the tracks in order to trace SPZs that were undetected by the program, or to delete those that were incorrectly detected as SPZ according to criterion of the operator. Specifically, we used object “Spots” in Imaris for the tracking of the SPZs. We set the estimated diameter of 8 *µ*m with background subtraction. We then set the filter type for the spots as ‘Quality’ that was dependent for each movie. The maximum distance (maximum search distance for an object to be considered part of the same track) was 10 *µ*m and the maximum gap size (maximum number of time frames that an object can go undetected and still be considered part of the same track) was 3. Finally, we used ‘Track Duration’ filter to select for tracks with a sufficient number of movements that was dependent on the movie (and operator). After all of these parameters are set and the initial automatic tracking is performed by Imaris, the operator then proceeded to modify the tracks by using Imaris manual corrections. We acknowledge that the manual editing of the tracks is probably operator-dependent. In total, we tracked 7,507 SPZs: 3,879 for Pf and and 3,628 for Py.

#### Cleaning trajectory data

As we have discovered previously^22,23^, when tracking cells with Imaris, its default settings allow several positions of a cell to be missing and still be considered as a single track; in our analyses this default value was 3. Such segmentation results in missing timeframes in the cell coordinate data that may lead to incorrectly calculated characteristics of a cell. Importantly, missing time frames in coordinate data exist in other published datasets in which movies were processed using other software^22,23^. We followed our previously outlined methodology to “clean” the trajectory data^22,23^. Specifically, for any time gap for a track with a given track ID that is greater than the imaging frequency of 1 sec, the track for that track ID is split into two tracks with unique track IDs before and after that time point. Such splits in track ID are repeated for all non-continuous time points. Cleaning of the data results in more trajectories that are typically shorter in duration. For example, for the data for Pf SPZs imaged 5 min after inoculation, we have 502 original tracks. Cleaning the data resulted in 1,191 tracks with 872 having 2 or more time points. We refer the original dataset as “unclean or original dataset” (**UD**) and the dataset resulting after splitting the tracks with missing time frames as “cleaned dataset” (**CD**).

#### Classifying SPZ trajectories as motile or immotile using maximal spatial spread

During initial inspection of the trajectory data, we observed that a considerable proportion of trajectories consisted of short, low-amplitude oscillations or “buzzing” movements that did not result in meaningful displacement. These trajectories often originated from sporozoites exhibiting local jitter or from segmentation and tracking artefacts near the detection threshold. Previous studies have used various methods to exclude trajectories that appear immotile (**Table 1**); in particular, for these data Hopp *et al*. ^13^ used overall displacement > 2 *µ*m to define motile trajectories. It is important to notice that using overall displacement (i.e., displacement from first to last position) to define motile SPZs may exclude circling parasites. We propose an alternative metric — maximal spatial spread *S* that is the maximal Euclidean displacement covered by a trajectory defined for parasites moving in 2D as

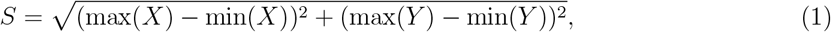

where *X* and *Y* denote the instantaneous coordinates of the sporozoite along the two-dimensional imaging plane. This maximal spatial spread can be easily extended to trajectories in 3D by adding calculation for Z coordinate in **eqn. (1)**. We use a threshold *S*_threshold_ for the maximal spatial spread to define a trajectory as motile (*S* > *S*_threshold_) or immotile (*S* ≤ *S*_threshold_). We implemented this filtering step in R as a custom function `filter tracks by spread()`, which executes the following sequence of operations:

1. calculates the per-track spread *f* using the maximum and minimum *X*–*Y* coordinates;
2. identifies all track IDs meeting both the spatial and point-count requirements;
3. retains only qualifying tracks and reports the number and percentage of removed trajectories; and
4. outputs a filtered dataset that preserves the structure of the original data table.

### Statistics

#### Quantifying mean squared displacement of SPZs

To characterize overall displacement of SPZs over time we calculated the mean squared displacement (**MSD**) as

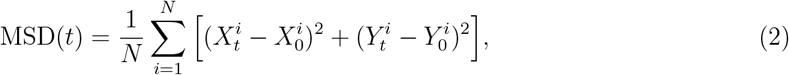

where *N* is the total number of tracks, 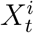 and 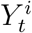 are the coordinates of the *i* track at time *t*, and 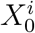 and 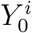 are the coordinates of the *i* track at the initial time *t* = 0. Note that we calculated MSD in 2D because our data only had one z-slice, and thus, all SPZs have identical z-coordinate. To calculate MSD we pooled the data from different experiment performed for the same SPZ species (Pf or Py) and measured at a defined time after SPZ inoculation (e.g., 5 min post-inoculation). For agents performing pure Brownian walks, MSD changes linearly with time (or similarly, log(MSD) ~ *γ* log *t* with *γ* = 1), and for agents performing correlated random walks (when movement velocity vectors are correlated), MSD initially increases faster than linearly with time (i.e., log(MSD) ~ *γ* log(*t*) with *γ* > 1)^22,23^. Slopes *γ* > 1 independently of the delay/time *t* typically indicates super-diffusive agents, e.g., Levy walkers^24–26^. We therefore tested how the rate of initial increase in MSD with time *γ* depends on the Plasmodium species, the time since inoculation, and *f*_threshold_ value used to define motile SPZs. To calculate MSD sloes *γ* we used data for the first 30 sec of displacement and performed linear regression of log(MSD) vs. log(*t*). When characterizing MSD curves we also calculate MSD value *µ* observed at *t* = 1 sec.

#### Quantifying distribution of turning angles for SPZs

Few studies typically calculate turning angles (**TAs**), the angle between two sequential movement vectors, for skin SPZs ^8,27,28^ even though he distribution of turning angles may inform about specific movement patterns^20,21^. Here we extend the methodology used to characterize movements of T cells in 3D, including movements biased in respect to an infection site^22,29,30^, to quantify different modes of SPZ movement in the skin. For that purpose we propose to use von Mises (**vM**) distribution that is a natural way to characterize biased angle distributions in 2D:

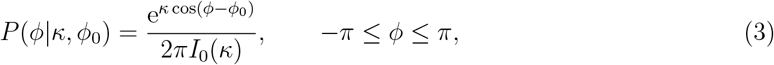

where *φ* and *φ*_0_ are the current (sample) and defined angles, *κ* is the concentration parameter and *I*_0_(*κ*) is the modified Bessel function of the first kind of order 0. Concentration parameter *κ* determines bias towards the defined angle *φ*_0_ with *κ* → 0 defining random/uniform (on circle) turning angles^31^. When calculating turning angles in 2D it is typical to ignore the sign of the angle and only consider the absolute value; in these cases 0 ≤ *φ* ≤ *π* and therefore strictly speaking, the vM distribution in **eqn. (3)** needs to be rescaled by a factor of 2. However, this rescaling typically does not impact the estimation of the concentration parameter *κ*. To fit the vM distribution to data we use likelihood approach, the negative log-likelihood (**NLL**) is defined as

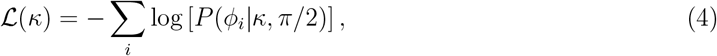

where *φ*_*i*_ are the measured TAs and the peak of the vM distribution is at *φ*_0_ = *π/*2. Best fit *κ* is found by maximizing L (**eqn. (4)**). Note that TA distribution of randomly moving agents in 2D is uniform between −*π* and *π* (i.e., when *κ* → 0 in **eqn. (3)**).

When analyzing SPZ turning angle distribution data we found that a single vM distribution typically does not accurately describe the data that often have peaks at 0 and 180 degrees^29,31^. Therefore, we use a mixture of vM distributions that are centered around *φ*_0_ = 0, 90 or 180 degrees with estimated weights *f*_*i*_; the NLL for a mixture of three vM distributions is

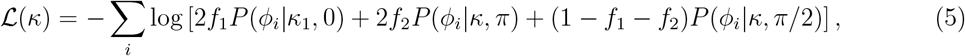

where *f*_1_ and *f*_2_ are the frequency of TAs that are biased towards 0 (moving forward) or 180 (turning back) degree, with the remaining fraction 1 − *f*_1_ − *f*_2_ denoting 90 degree turns (may indicate circling). It is also possible to fit 2 vM distributions to TA data assuming bias only towards forward direction (*φ*_0_ = 0) or reverse direction (*φ*_0_ = *π*). The quality of the fit of a single vs. mixture of vM distributions is assessed using likelihood ratio test^32^.

#### Defining SPZ movement modes using moveHMM library in R

To complement the analysis of SPZ movement patterns based on TA distribution (**eqn. (3)**) and to quantify potential SPZ movement types, we used a recently proposed method, based on hidden Markov models (**HMMs**) and implemented in the `moveHMM` package in R. moveHMM characterizes movement of agents via several independent “states” or modes and allows to estimate parameters of these states/programs using information on the joint distribution of turning angles and step lengths^20,21^. For this analysis we used cleaned datasets (clean track IDs, time in seconds and two-dimensional positions in *µ*m). In our preliminary analyses we found that fitting HMMs to full dataset without removing any potentially immotile trajectories (thus, assuming *f*_threshold_ = 0), often did not converge or did not result in biologically meaningful parameter estimates. Therefore, in our main analyses with moveHMM we discarded tracks 1) with fewer than three positions, 2) with maximal spatial spread *S* < *S*_threshold_ = 5 *µ*m (**eqn. (1)**). To generate the curated data we used `prepData` function in moveHMM.

We then fit HMMs assuming two or three different movement states to the data; we used gamma distribution for step lengths and von Mises distribution (**eqn. (3)**) for turning angles when fitting the models to data. We used initial guesses for step-length means from empirical quantiles of the observed distribution (25th–75th percentiles for the 2-state model and 20th–50th—-80th percentiles for the 3-state model), with standard deviations set to half of the corresponding means. We used initial guesses for bias in angles to reflect biologically plausible biases: the mean directions were initialized at *π* (≈ 3.14 rad), 0, and 0 for the three states, representing reversal, forward persistence, and near-random turning, respectively. The concentration parameters (*κ*) were scaled from the pooled circular statistics of the observed angles, with *κ*_1_ < *κ*_2_ < *κ*_3_ to reflect a hierarchy from diffuse to strongly persistent movement. The calibration quality was checked by comparing the theoretical quantiles of the fitted gamma distributions with the sample quantiles of the empirical step-length distribution to ensure stable initialization and convergence. We evaluated the alternative models (e.g., 2 vs. 3 states) using AIC, and most-likely state sequences were obtained using the Viterbi algorithm ^20,33^. We used best parameter estimates to simulate SPZ movement with routine `SimData`. All analyses were performed in R (version 4.4.2) with defined random seeds to ensure reproducibility.

#### Statistical significance and effect size

To compare averages (e.g., average displacement) we used unpaired Student’s *t*-test with p-values less than or equal to 0.05 being statistically significant; p-values smaller than 0.001 were reported as *p* < 0.001 due to their exceedingly small magnitude, rather than displaying the exact values. To measure the magnitude of the effect size between the different populations across all time points post-inoculation and between the different species, we calculated Cohen’s *d* ^34^. The interpretation of the values obtained for Cohen’s *d* for effect size are indicated as follows:

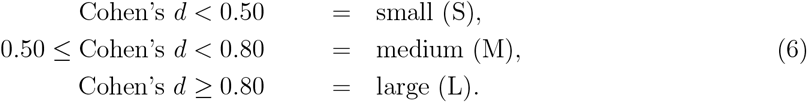

To evaluate the magnitude of the effect size of the linear regression models, we calculated Cohen’s *f*^2 34,35^; it can be interpreted as follows:

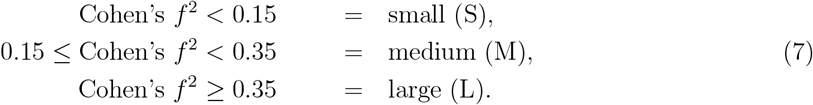

## Results

### Classifying SPZ trajectories as motile or immotile is not always straight-forward

Our review of the previously published studies suggested a variety of metrics used to define motile SPZs (**Table 1**); however, the choice of the metric and specific threshold value used to define SPZs as motile has been rarely discussed. For example, metric defining motile SPZs when they are circling requires defining “circling”; this may be visually straightforward for few parasite but may be mathematically challenging when using automatic classifiers for hundreds of trajectories ^36^. We propose a new metric, maximal spatial spread (**MSS** or *S*, **eqn. (1)**), to characterize overall SPZ motility; this metric takes into account all positions of a given SPZ during the observational period and is the maximal Euclidean distance between minimum and maximum positions taken by the parasite (**Figure 2**A). We calculated *f* for every cleaned SPZ trajectory in our newly generated dataset of 22 movies on movement of Pf and Py SPZs in murine skin at different times after syringe inoculation into the skin (see Materials and methods for detail, **Figure 2**B-C and **Movies 1–2**). We found very wide distribution of MSS ranging from 0.1 *µ*m to nearly 10^2^ *µ*m that was persistent with time since inoculation. Unfortunately, there was no clear division of the distribution data into parasites with small and large MSS for Pf SPZs (**Figure 2**B); however, for Py SPZs the distribution could be divided into SPZs with small (*f* < 5 *µ*m) or large (*f* > 5 *µ*m) maximal spatial spread (**Figure 2**C). Interestingly, the proportion of parasites higher MSS declined with time since inoculation suggesting reduction in the frequency of more motile SPZs with time. This analysis suggests that depending on the dataset there may or may not be a clear separation in the MSS that would unambiguously classify SPZs as motile or immotile and thus, the decision of assigning parasites as motile may in some cases be semi-arbitrary.

**Figure 2:**
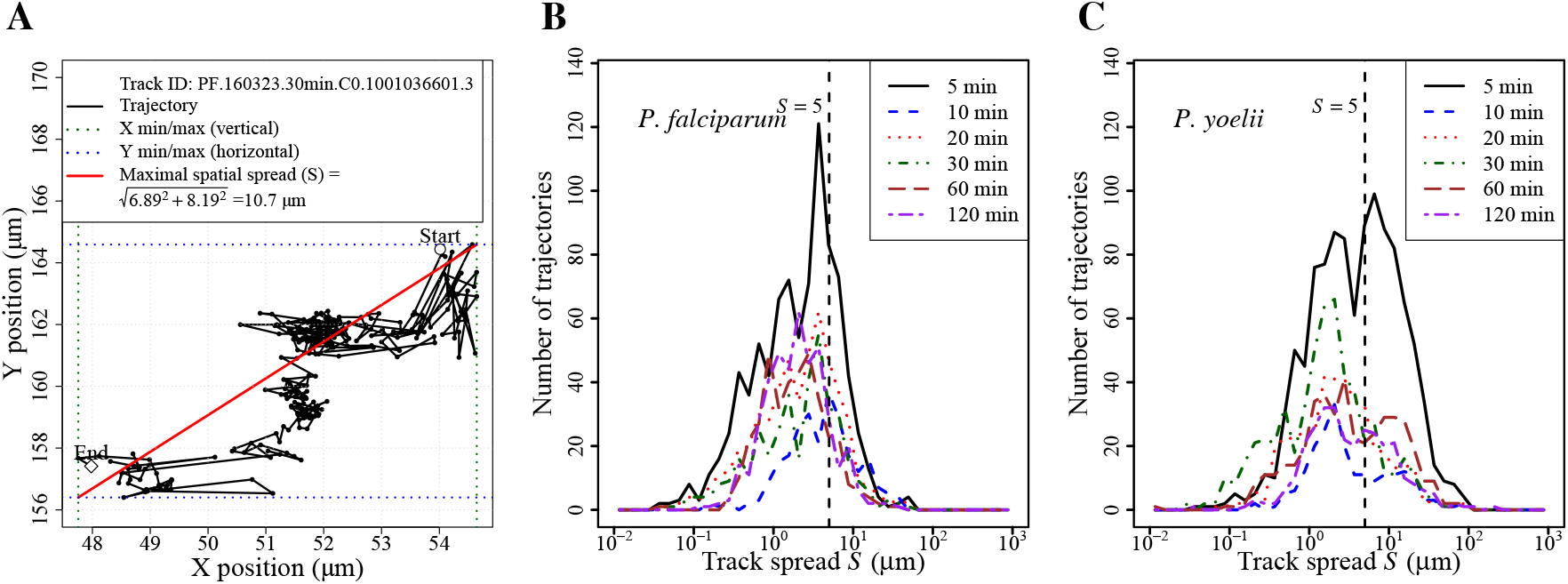
Plasmodium sporozoites exhibit a wide range of maximal spatial spread. We calculated the maximal spatial spread for every trajectory of Pf and Py SPZs in data from our newly processed 22 movies. **A**: Representative trajectory of a Pf SPZ (track ID PF.160323, 30 min post-inoculation) illustrating the calculation of maximal spatial spread *S* (see **eqn. (1)**). Vertical and horizontal dashed lines denote the minimum and maximum *x*- and *y*-coordinates of the trajectory. (**B**–**C**) Distributions of maximal spatial spread *S* for all tracked Pf (**B**) or Py (**C**) SPZs at different times post-inoculation (5–120 min, denoted by lines of different colors and styles). In B-C, vertical dashed lines denote *S*_threshold_ = 5 *µ*m.

We next explored how changing threshold value for the maximal spatial spread *S*_threshold_ would influence basic movement characteristics (**Supplemental Table S1** and **Table 2**). As expected, increasing *S*_threshold_ naturally led to fewer trajectories classified as motile for both Pf and Py SPZs and was relatively independent of the time since SPZ inoculation (**Supplemental Table S1**). Yet, the fraction of motile SPZ was highly variable between different datasets ranging from 3% of Pf SPZs at 120min after inoculation to 54% of Py SPZs at 120min post inoculation. Interestingly, there was no obvious trend in change of the fraction of motile SPZs with time since inoculation independently of the *S*_threshold_ values (**Supplemental Table S1**). Increasing *S*_threshold_ did not influence the mean track duration (i.e., did not select for longer or shorter tracks) but did result in larger final displacements, larger average instantaneous or average track speeds, or in the slope *γ* of how mean squared displacement (**MSD**) changes with time (**Table 2** and **Supplemental Figure S1**). Interestingly, for the same *S*_threshold_ value, Py SPZs exhibited larger values for all of the movement characteristics (final displacement, instantaneous or mean speeds, and slope *γ*) as compared to Pf SPZs. Taken together, our analysis shows that estimates of basic motility parameters of Plasmodium SPZs strongly depend on the value of MSS threshold used to classify tracks as motile.

**Table 2:** Increasing the maximal spatial spread threshold *S*_threshold_ results in larger final displacements, higher speeds, larger early MSD slopes for both *P. falciparum* and *P. yoelii* SPZs. We show the average final displacement, instantaneous speed, mean speed, mean early MSD slope (*γ*), and mean track duration calculated for each maximal spatial spread threshold (*S*_threshold_ = 2–10 *µ*m). We calculated these values by using all tracks for Pf or Py SPZs for all times after the inoculation.

| Species | Spatial spread threshold<br>$S_{\text{threshold}}$ ($\mu\text{m}$) | Final Displacement<br>($\mu\text{m}$) | Instantaneous Speed<br>($\mu\text{m/s}$) | Mean Speed<br>($\mu\text{m/s}$) | Mean MSD Slope<br>$\gamma$ | Mean Track Duration<br>(s) |
| --- | --- | --- | --- | --- | --- | --- |
| *P. falciparum* | 2 | 4.32 | 0.344 | 0.484 | 0.93 | 122.2 |
|  | 5 | 7.89 | 0.403 | 0.498 | 1.07 | 132.2 |
|  | 7 | 10.9 | 0.429 | 0.522 | 1.22 | 128.4 |
|  | 10 | 15.6 | 0.449 | 0.542 | 1.36 | 128.4 |
| *P. yoelii* | 2 | 7.23 | 0.639 | 1.15 | 0.967 | 63.3 |
|  | 5 | 11.0 | 0.781 | 1.28 | 0.981 | 59.6 |
|  | 7 | 13.6 | 0.852 | 1.26 | 1.00 | 60.2 |
|  | 10 | 16.6 | 0.903 | 1.22 | 1.05 | 59.8 |

### Selection criteria for motile/immotile tracks influences change in movement characteristics with time since inoculation

Previous analyses of trajectory data of Pf and Py SPZs found that the parasites do not differ significantly in some of their movement characteristics, e.g., final displacement or average speeds, and found that average SPZ speed tends to decline with time since SPZ inoculation into the skin for Pf and Py SPZs^17^. To confirm these observations while using of novel methodology to select for motile SPZs, we compared how final displacement and average speeds of Pf and Py SPZs change with time since inoculation for 22 movies we had re-analyzed. In these experiments, we processed movies for Pf and Py SPZs tracked for 4 minutes 5, 10, 20, 30, 60, and 120 min following deposition into the dermis (**Figure 3**). To simplify the analysis we either selected all tracks for the analysis (*S*_threshold_ = 0) or used intermediate value (*S*_threshold_ = 5 *µ*m) to define motile SPZs (**Figure 2**C).

**Figure 3:**
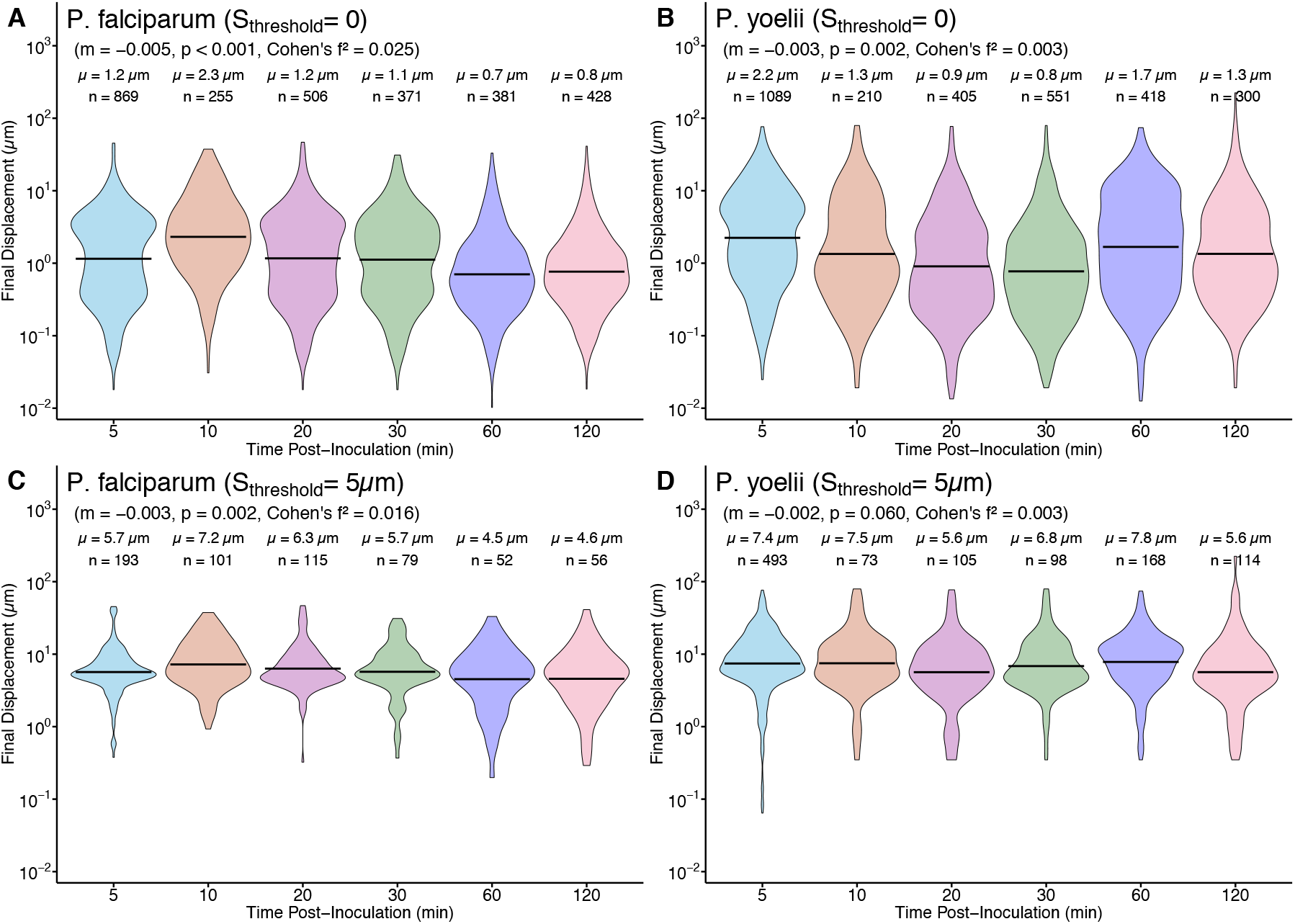
Larger values of maximal spatial spread threshold *S*_threshold_ leads to higher final displacement of SPZs in skin. We quantified the final displacement (displacement between 1st and last track position) of *P. falciparum* (A,C) and *P. yoelii* (B,D) sporozoites at multiple time points post inoculation. We performed these analyses with a cleaned dataset that has been corrected for missing time frames in cell coordinate data (see Materials and methods for detail and **Supplemental Figure S2** for similar analysis with original/unclean data). Panels A–B are for all trajectories (*S*_threshold_ = 0), whereas panels C–D include only trajectories with *S* > *S*_threshold_ = 5 *µ*m of maximal spatial spread. We visualized displacement distributions using violin plots with geometric mean (*µ*) and sample size (*n*) indicated on individual plots. We performed linear regression to estimate slope (*m*), associated *p* values, and Cohen’s *f*^2^ (**eqn. (7)**) sizes for change in final displacement with time since inoculation.

At *S*_threshold_ = 0, Pf SPZs exhibited relatively modest displacement values (geometric means of 1.2–0.8 µm) that gradually decreased over time (slope *m* = − 0.005, *p* = 9.8 × 10^*−*17^; Cohen’s *f*^2^ = 0.025, **Figure 3**A). *P. yoelii* showed higher initial displacements (2.2 µm at 5 min) but a similar temporal decay pattern (*m* = −0.003, *p* = 0.002; *f*^2^ = 0.003), indicating a general reduction in motility intensity over time for both species if we assume that all SPZ in our dataset are motile (**Figure 3**B).

In contrast, using a moderate MSS threshold (*S*_threshold_ = 5 *µ*m) to define motile SPZs markedly reshaped the final displacement distributions. This filtering step for Pf SPZs resulted in geometric mean final displacements of 5.7–4.6 µm and a shallower decay slope (*m* = −0.003, *p* = 0.002; *f*^2^ = 0.016, **Figure 3**C). Similarly, Py SPZs exhibited consistently higher displacements under the same *S*_threshold_ (7.4–5.6 µm) with minimal temporal decay (*m* = − 0.002, *p* = 0.06; *f*^2^ = 0.003, **Figure 3**D).

The impact of higher threshold value was strongest at early time points (5–20 min after SPZ inoculation), where trajectories with very small MSS comprised a larger fraction of the population (**Supplemental Table S1**). By removing these low-dispersion tracks, we found clearer species-level differences: Py SPZs maintained larger displacements and weaker temporal decline in final displacement than Pf SPZs. The results were similar if we use original/uncleaned trajectory data for the analysis where higher *S*_threshold_ values resulted in larger displacement and smaller change in final displacement with time since SPZ inoculation (**Supplemental Figure S2**).

We performed similar analyses when calculating the average speed per trajectory (**Figure 4**). When classifying all trajectories as motile (*S*_threshold_ = 0), Pf SPZs displayed a gradual decline in mean speed over the course of the 120 min observation period (**Figure 4**A). Mean speeds decreased slightly from 0.35 µm/s at 5 min to 0.31 µm/s at 120 min (sample sizes: *n* = 872 and *n* = 428, respectively). Linear regression confirmed a significant negative temporal trend (slope *m* = − 0.002, *p* < 0.001, Cohen’s *f*^2^ = 0.014). In contrast, Py SPZs exhibited consistently higher mean speeds at all time points (**Figure 4**B). Geometric means were 0.86 µm/s at 5 min and 0.49 µm/s at 120 min (*n* = 1089 and *n* = 301). Although a decline was also observed over time, the decline slope was larger than that for Pf SPZs (*m* = −0.004, *p* < 0.001, Cohen’s *f*^2^ = 0.020, **Figure 4**B).

**Figure 4:**
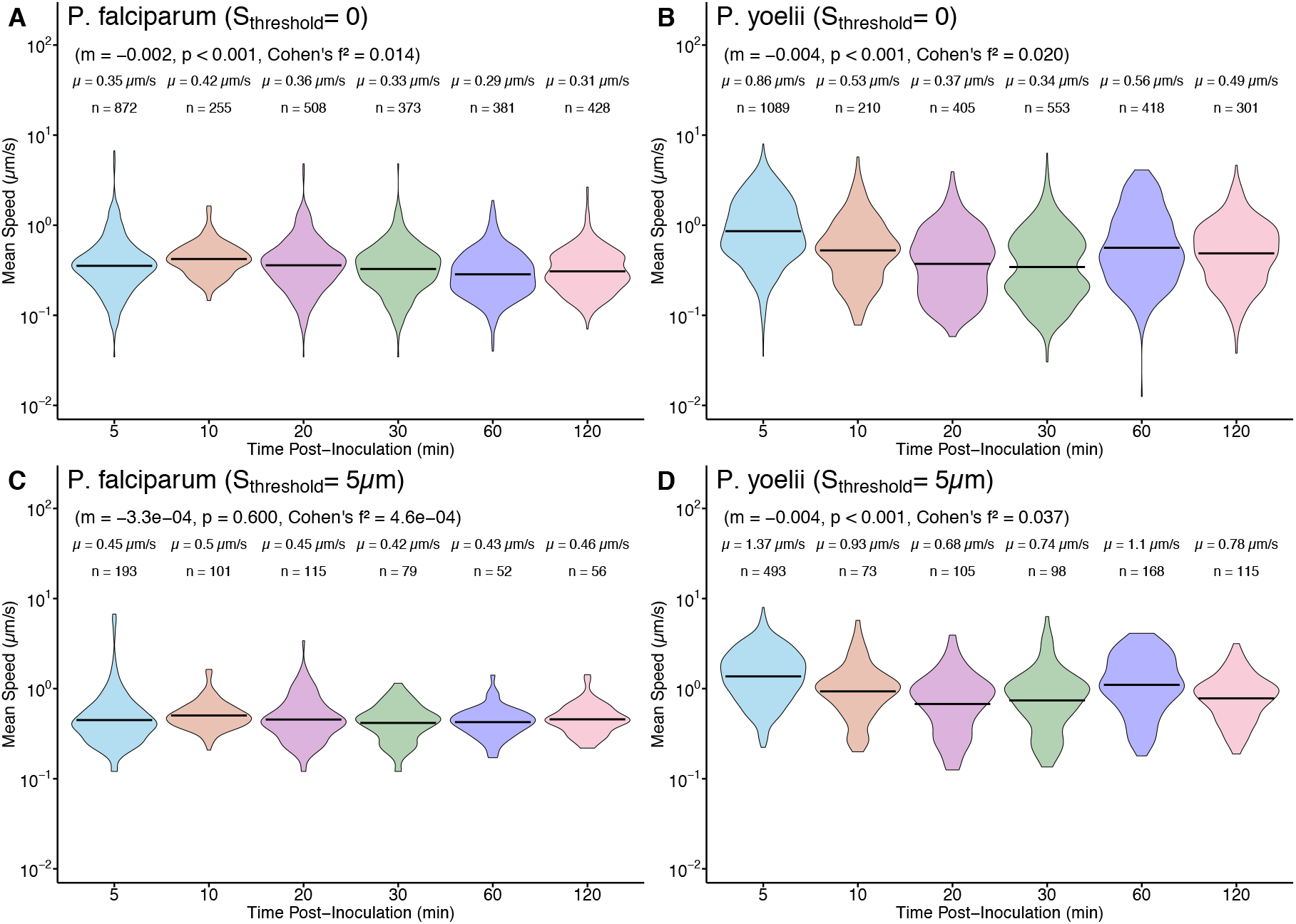
Larger maximal spatial spread threshold values lead to higher average speed of SPZs. We computed mean speeds of Pf (A,C) or Py (B,D) SPZs at multiple time points post inoculation. We used clean dataset for the analysis (see **Figure 3** for other details and **Supplemental Figure S3** for analysis of unclean data). Panels A–B are for all trajectories (*S*_threshold_ = 0), and panels C–D are for trajectories with maximal spatial spread *S* ≥ *S*_threshold_ = 5 *µ*m. We also show the average speed for all tracks *µ* and the number of tracks *n* per dataset. We applied linear regression to estimate slope (*m*), *p* values, and Cohen’s *f*^2^.

When motile tracks were defined with MSS threshold *S*_threshold_ = 5 *µ*m we found large changes in average speeds for both species, particularly at early time points (**Figure 4**C–D). Selection for SPZ with larger MSS elevated mean speeds at all time points (0.45–0.46 µm/s) for Pf SPZs and flattened the temporal trend (*m* = − 0.00033, *p* = 0.6, *f*^2^ = 0.00046). This indicates that the initial negative slope was largely driven by the inclusion of numerous low-spread trajectories at early post-inoculation intervals. Py SPZs in the selected dataset retained higher mean speeds with values ranging from 1.37 µm/s at 5 min to 0.78 µm/s at 120 min. Although a decline remained detectable (*m* = − 0.004, *p* < 0.001, *f*^2^ = 0.037), the decline slope was smaller further suggesting that change in average speed of SPZs with time may depend on the methodology used to define motile SPZs. Py SPZs exhibited higher overall motility across several tested threshold values, with mean speeds ranging from 1.15 to 1.28 *µ*m/s and displacements from 7.23 to 16.6 *µ*m (**Table 2**). Results were similar for unclean data (**Supplemental Figure S3**). Together, these results demonstrate that the observed temporal dynamics of final displacement and mean speed depend strongly on which SPZs are deemed motile.

### Larger maximal spatial spread thresholds selects for trajectories displaying persistent movement

We next analyzed how the mean squared displacement (**MSD, eqn. (2)**) of Pf and Py SPZs changes with time; this allows to evaluate how SPZs may be dispersing from the inoculation site and how evaluation of the dispersal may depend on deciding which trajectories are motile (**Figure 5** and **Table 3**). MSD curves are one of the metrics allowing to distinguish normal (Brownian) diffusion from other types of walks, e.g., Levy walks^22,25^.

**Table 3:** Increasing the spatial spread threshold results in larger MSD slopes. We calculated the short-time MSD slopes (*γ*) for *P. falciparum* and *P. yoelii* across multiple post-inoculation time points at different values of spatial spread threshold *S*_threshold_. For each threshold value, we computed the MSD slope for trajectories with *S* > *S*_threshold_ and estimated slope *γ* by using linear regression for log(MSD) versus log(*t*) over the first 30 s of movement.

| Spatial spread threshold<br>($S_{\text{threshold}}$, $\mu\text{m}$) | Time Post Inoculation<br>(min) | MSD slope ($\gamma$) | |
| --- | --- | --- | --- |
|  |  | *P. falciparum* | *P. yoelii* |
| 0 | 5 | 1.04 | 0.94 |
|  | 10 | 1.05 | 1.03 |
|  | 20 | 1.00 | 1.05 |
|  | 30 | 1.03 | 1.11 |
|  | 60 | 0.81 | 0.94 |
|  | 120 | 0.91 | 0.99 |
| 2 | 5 | 0.96 | 0.92 |
|  | 10 | 1.03 | 1.04 |
|  | 20 | 0.98 | 0.94 |
|  | 30 | 0.96 | 0.91 |
|  | 60 | 0.89 | 0.94 |
|  | 120 | 0.90 | 0.99 |
| 5 | 5 | 1.07 | 0.92 |
|  | 10 | 1.09 | 1.07 |
|  | 20 | 0.99 | 1.07 |
|  | 30 | 1.09 | 0.96 |
|  | 60 | 1.08 | 1.02 |
|  | 120 | 1.19 | 1.02 |
| 7 | 5 | 1.31 | 0.94 |
|  | 10 | 1.26 | 1.12 |
|  | 20 | 1.08 | 1.15 |
|  | 30 | 1.21 | 1.11 |
|  | 60 | 1.14 | 0.97 |
|  | 120 | 1.22 | 1.07 |
| 10 | 5 | 1.42 | 0.98 |
|  | 10 | 1.39 | 1.13 |
|  | 20 | 1.29 | 1.19 |
|  | 30 | 1.30 | 1.17 |
|  | 60 | 1.23 | 1.01 |
|  | 120 | 1.43 | 1.14 |

**Figure 5:**
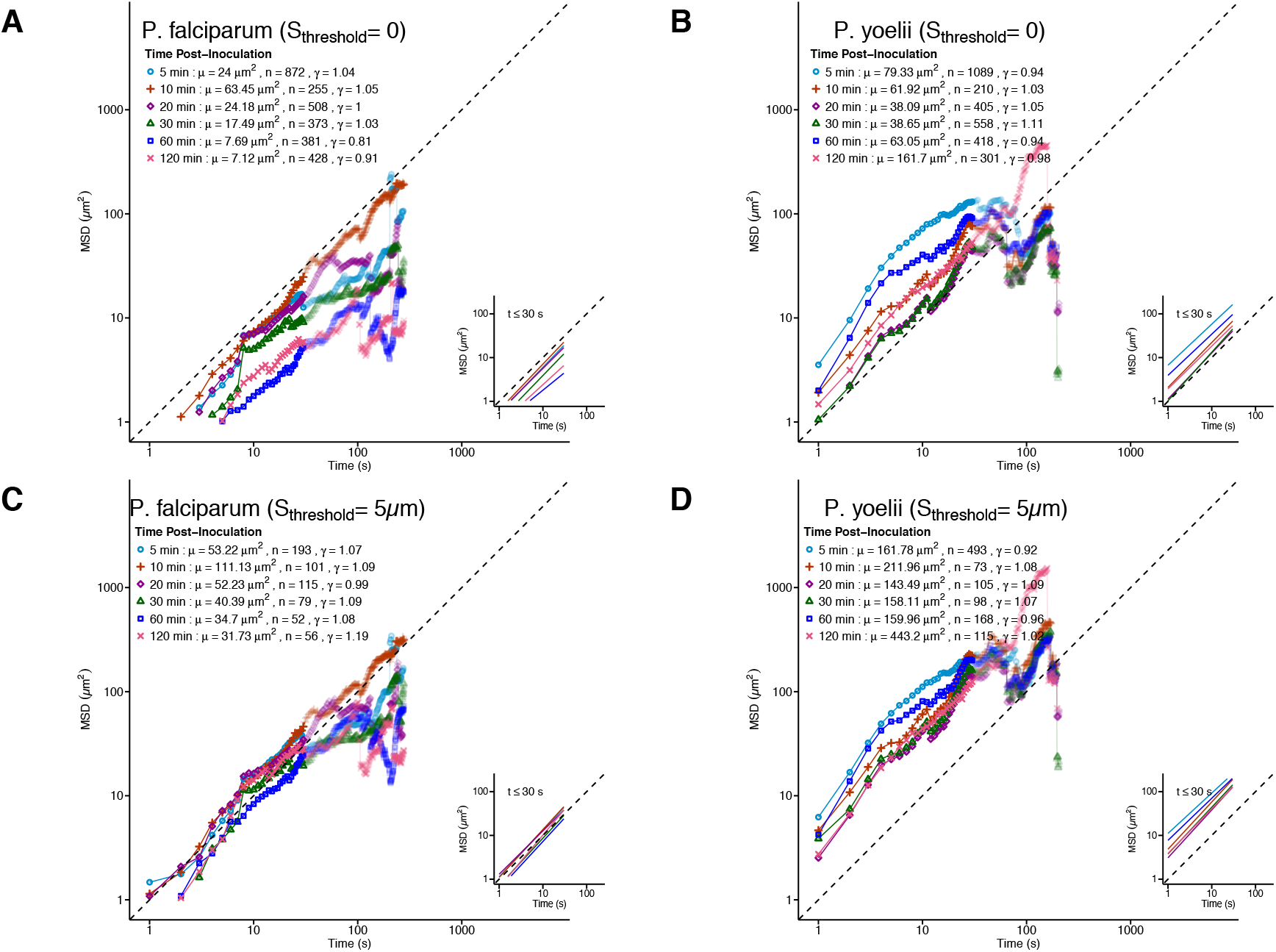
Displacement-based trajectory selection increases inferred motility persistence. We calculated MSD curves for *P. falciparum* (A,C) and *P. yoelii* (B,D) trajectories across time post inoculation. Panels A–B are for all trajectories (*S*_threshold_ = 0), whereas panels C–D are for trajectories with *S* ≥ *S*_threshold_ = 5 *µ*m (see **eqn. (1)**). We estimated the short-time slope (*γ*) of log(MSD) versus log(*t*) over the first 30 s (insets). MSD slopes for other values of *S*_threshold_ are listed in **Table 3**. In the panels we show the MSD value at 1 sec (*µ*), number of trajectories (*n*), and estimated MSD slope (*γ*).

When including all trajectories in the analysis (*S*_threshold_ = 0), Pf SPZs displayed moderate early dispersal at 5–20 min post inoculation, with MSD slopes (*γ*) in first 30 sec around 1.0–1.05 (e.g., 5 min: MSD value at *t* = 1 sec *µ* = 24 *µ*m^2^, *γ* = 1.04; 10 min: *µ* = 63.45 *µ*m^2^, *γ* = 1.05, **Figure 5**A and **Table 3**). As time progressed, both magnitude and slopes declined (e.g., 60 min: *µ* = 7.69 *µ*m^2^, *γ* = 0.81), indicating increasing prevalence of weakly displacing parasites. Py SPZs showed higher MSD magnitudes and relatively stable slopes around unity at early time points (e.g., 5 min: *µ* = 79.33 *µ*m^2^, *γ* = 0.94; 20 min: *µ* = 38.09 *µ*m^2^, *γ* = 1.05), with modest variation over time, consistent with faster but more stochastic migration compared to Pf SPZs (**Figure 5**B and **Table 3**). Overall, change in MSD with time was close to linear, consistent with previous observations ^13^.

Selecting trajectories with *f* > *S*_threshold_ = 5 *µ*m resulted in substantially increased MSD values and MSD increase slopes across time points in both species (**Figure 5**C-D and **Table 3**). For *P. falciparum*, mean MSD at 5 min rose to 53.2 µm^2^ with *γ* = 1.07, and remained above 30 µm^2^ with slopes close to or above 1.0 even at later time points (e.g., 120 min: *µ* = 31.73 *µ*m^2^, *γ* = 1.19) indicating super-diffusive displacement. This flattening of temporal decay and increase in slope indicates that removing trajectories with small MSS enriches for directionally persistent tracks.

Py SPZs with *f* > *S*_threshold_ = 5 *µ*m exhibited even higher MSD values post-filtering (e.g., 5 min: *µ* = 161.78 *µ*m^2^, *γ* = 0.92; 10 min: *µ* = 211.96 *µ*m^2^, *γ* = 1.08; 120 min: *µ* = 443.2 *µ*m^2^, *γ* = 1.02), with even higher initial slopes *γ* in the first 10-15sec of imaging (**Figure 5**D). Together, these results demonstrate that MSD-based characterization of SPZ movement is sensitive to the decision which trajectories are defined as motile (but also on deciding the time period for calculating the slope *γ*).

Furthermore, Pf and Py SPZs do differ in their motility as Pf SPZs display more pure Brownian-like motion while Py SPZs do show faster than linear initial displacement in the first 10-15 sec consistent with correlated random walks (**Figure 5**C-D).

### Larger maximal spatial spread threshold values result in significant changes in distribution of turning angles

To complement MSD analysis, we quantified turning angle distributions as a measure of directional persistence and reorientation behavior over time. High mean turning angles (**TAs**) near 90–120° indicate frequent reorientation and diffusive behavior, whereas lower mean TAs suggest persistent migration^22,37^. Interestingly, when analyzing all trajectories (i.e., *S*_threshold_ = 0), TAs for Pf SPZs were broadly stable over time, with mean angles between 109^0^ and 116^0^ across all time points suggesting frequent turning back (e.g., 5 min: *µ* = 113^0^; 60 min: *µ* = 114^0^; 120 min: *µ* = 115^0^, **Figure 6**A). Linear regression confirmed no significant temporal trend (*m* = 0.03, *p* = 0.26), indicating that although final displacement declined over time, turning angle characteristics of the population remained constant. Py SPZs showed slightly lower TAs at early time points (e.g., 5 min: *µ* = 96^0^, **Figure 6**B), but overall remained near 100^0^ with no significant change with time since inoculation (*m* = 0.023, *p* = 0.66). Overall, this pattern reflects a motility mode characterized by fast but frequent reorientation, consistent with initial MSD slopes close to 1.

**Figure 6:**
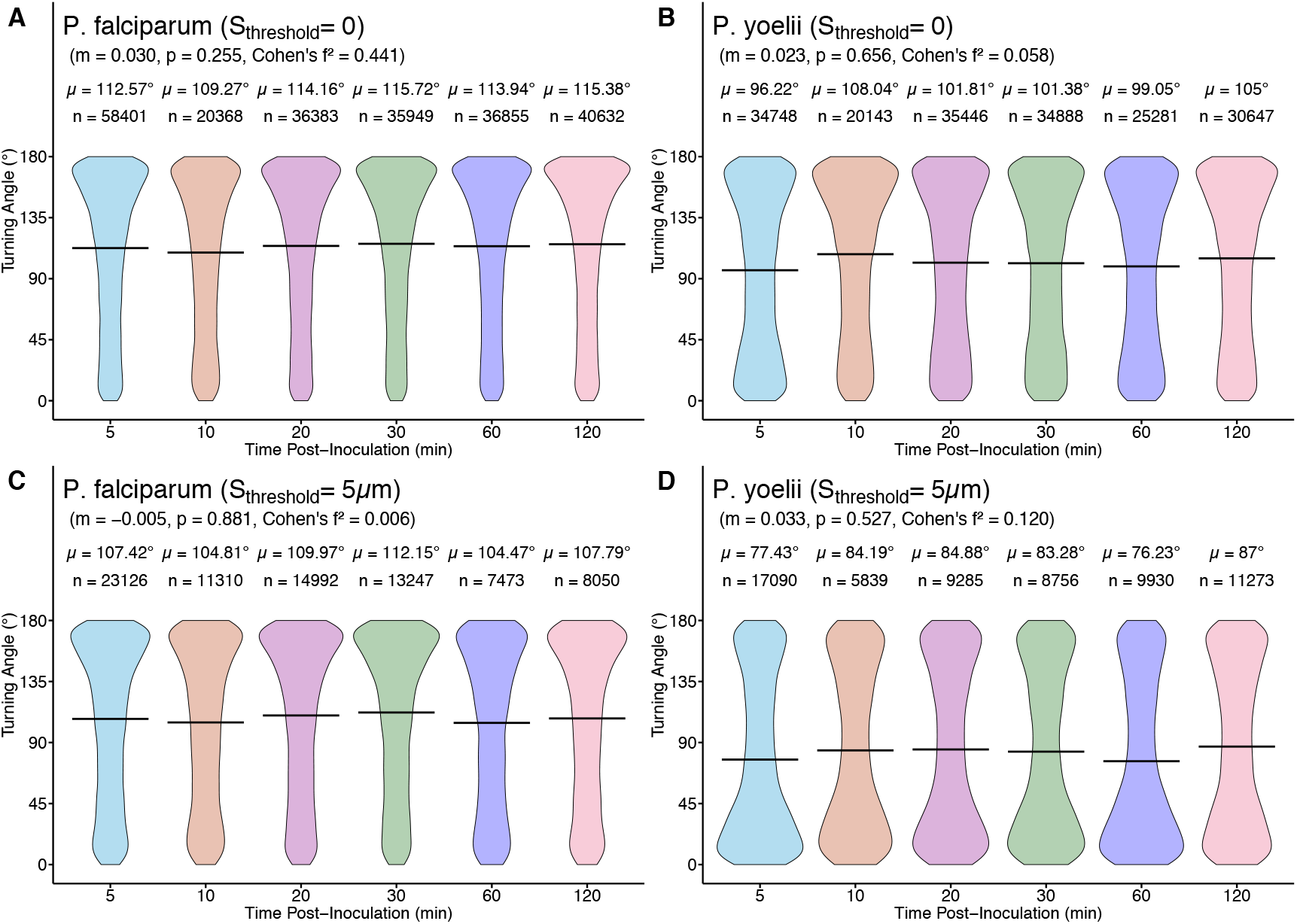
Larger spatial spread threshold values change distribution of turning angles for Py SPZs but not for Pf SPZs. We calculated turning angles for *P. falciparum* (A, C) and *P. yoelii* (B, D) trajectories at multiple time points post inoculation. Panels A–B are for all trajectories (*S*_threshold_ = 0), and panels C–D are for trajectories with *S* > *S*_threshold_ = 5 *µ*m (see **eqn. (1)**). Violin plots depict angle distributions with *µ* and *n* indicated. We applied linear regression fits (dashed lines) to estimate slope (*γ*), *p* values, and Cohen’s *f*^2^.

Importantly, selecting trajectories with *S* > *S*_threshold_ = 5 *µ*m as motile reduced mean TAs for both species (**Figure 6**C-D), indicating a selective enrichment of trajectories with persistent movement. For Pf SPZs, mean TAs stabilized around 105–112^0^, with no significant temporal trend (*m* = −0.005, *p* = 0.88). In contrast, for Py SPZs, mean TA decreased more substantially (e.g., 5 min: *µ* = 77^0^; 120 min: *µ* = 87^0^), yet with no significant temporal trend (*m* = 0.033, *p* = 0.53). Sub-selection of trajectories disproportionately affected TAs of Py SPZs further demonstrating that the methodology of deciding which trajectories are motile may influence inferred movement patterns (**Supplemental Figure S4**).

With sub-selection of trajectories with larger MSS, the divergence between the two species became clearer: Pf SPZs maintained moderate TAs with short persistence, whereas Py SPZs exhibited lower TAs indicating more persistent directional movement. Sub-selection of trajectories with higher *S*_threshold_ values resulted in fewer trajectories defined as motile (**Supplemental Table S1**). For *P. falciparum* with *S*_threshold_ = 2 *µ*m, approximately 40% of trajectories were defined as motile during early post-inoculation time points (5–60 min), but fewer than 10–15% were defined motile once the threshold exceeded 7–10 *µ*m (**Supplemental Table S1**). In contrast, 50–55% of Py SPZs were classified as motile at *S*_threshold_ = 2 *µ*m and 15–20% were still classified as motile at *S*_threshold_ = 10 *µ*m. The magnitude of reduction in motile fractions directly quantifies how subselection by maximal spatial spread modifies the apparent ensemble behavior: stringent thresholds remove weakly motile or stationary parasites, yielding datasets dominated by persistent, long-range gliders. Overall, this analysis revealed that the apparent motility behavior of the parasite population is sensitive to the criterion used for trajectory inclusion. Stricter filters progressively shift the aggregate behavior from mixed (diffusive + stationary) toward a more coherent, directionally persistent regime.

We have previously proposed methodology, based on von Mises-Fisher distribution, to characterize bias in T cell movement searching for Plasmodium parasites in the liver^30^. Here we have extended this methodology to quantify distribution of TAs of SPZs moving in 2D. The method is based on fitting a combination of 2 or 3 von Mises distribution to the turning angle distribution data; these distributions quantify the proportion of forward movements, reverse movements, and movement at 90 degrees (**eqn. (3)** and see Materials and methods for detail).

We found that typically at least 3 vM distributions are required to accurately describe the TA distribution (**Figure 7**). Interestingly, when considering all trajectories as motile (*S*_threshold_ = 0), TA distribution was dominated by reversal movements with relatively high concentration parameter *κ*_2_ with forward movements constituting only 30-40% (**Figure 7**A-B). Very few movements represented potential circling, defined as movements with TA=90^0^ (*f*_3_ = 10%). In contrast, sub-selecting trajectories with *S* > *S*_threshold_ = 5 *µ*m as motile changed the inferred movement pattern for Py SPZs but not for Pf SPZs (**Figure 7**C-D). Specifically, for Py SPZs now the majority of movements (*f*_1_ = 55%) are now in the forward direction at higher persistence (*κ*_1_ = 2.53). This is consistent with the super-diffusive displacement of Py SPZs observed for initial MSD change with time for *S*_threshold_ = 5 *µ*m (**Figure 5**D). Yet, there was limited change in vM distribution fits to TA data for Pf SPZs with predominance of reverse displacements (**Figure 7**C). Thus, the choice of criteria for selecting SPZ trajectories as motile influences changes in MSD and TA distributions, and thus, inference of the movement modes of SPZs in vivo.

**Figure 7:**
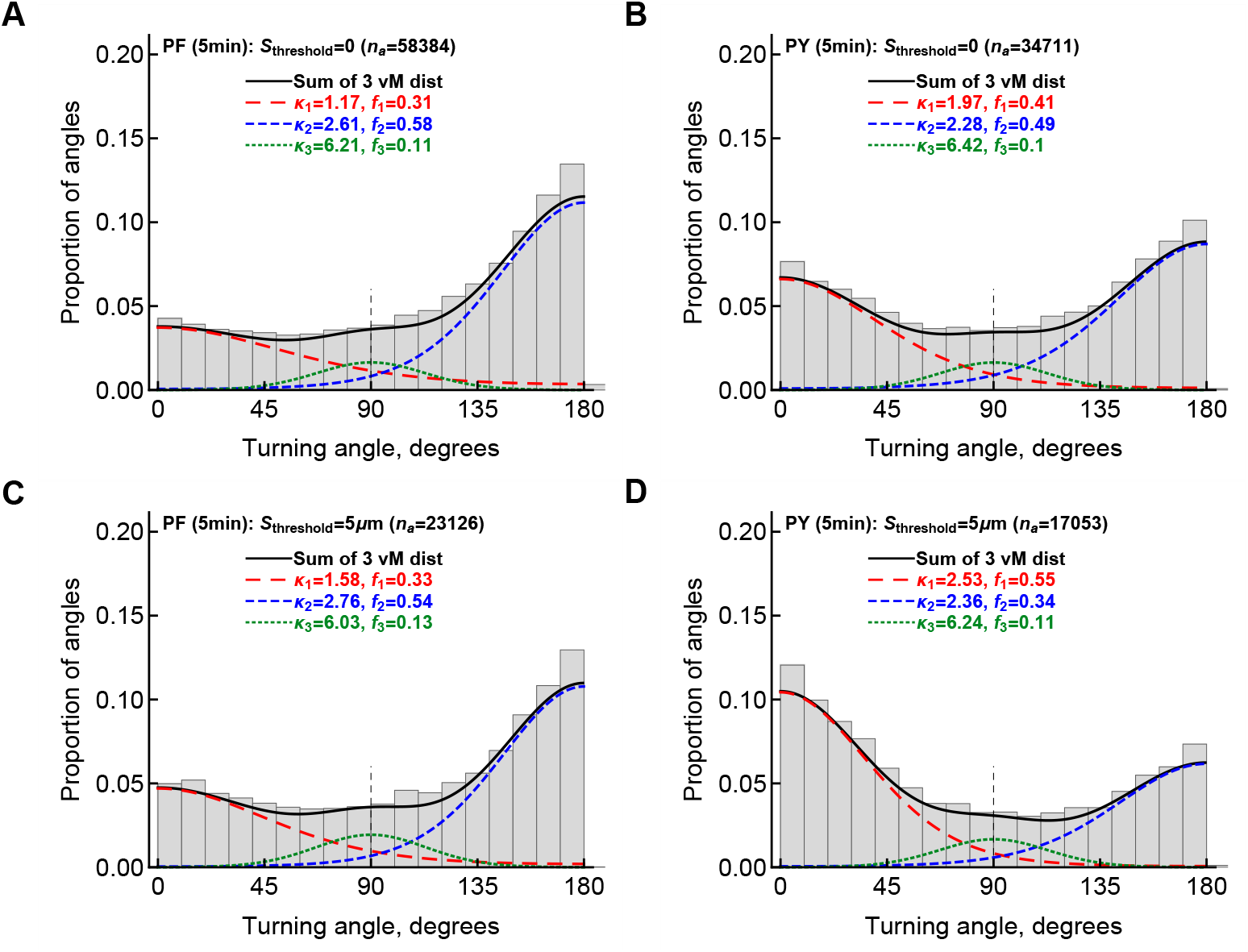
Larger maximal spatial spread threshold selects for persistently moving Py SPZs. We fitted the mixture of three vM distributions (**eqn. (3)**) to the turning angle (**TA**) data for Pf (**A&C**) or Py (**B&D**) sporozoites moving in the skin at 5 min after the inoculation by either using all data (*S*_threshold_ = 0, **A**-**B**) or excluding tracks with smaller than *S*_threshold_ = 5 *µ*m maximal spatial spread(**C**-**D**). The number of TAs (*n*_*a*_) is shown on individual panels. We show the proportion of forward movements, characterized by the concentration parameter *κ*_1_, as *f*_1_ (red dashed lines), the proportion of reversal movements as *f*_2_ (with *κ*_2_, blue small dashed lines), the proportion of right angle turns as *f*_3_ (with *κ*_3_, green dotted lines), and the sum of the three distributions (black continuous lines). Fits of a model consisting of one or mixture of two vM distributions were of lower quality than those presented on the panels (LRT, *p* < 0.01).

### Cleaning/splitting tracks does not impact average speed of the tracks in our datasets

We have previously shown that semi-automatic processing of imaging data generates T cell tracks that often have missing time frames; such missing time frames may in some cases result in misclassification of T cell movement program^22,23^. After generating the SPZ trajectory data we also found missing time frames for some trajectories; for most of our analyses we used cleaned data that split trajectories to generate novel trajectories that have sequential measurements of cell positions (see Materials and methods for detail). Because cleaning cell trajectories may influence various motility parameters, we tested if cleaning trajectories would change average trajectory speed. Interestingly, in contrast with our results for T cells, cleaning SPZ tracks did not influence the average speeds when taking all trajectories into account (i.e., with *S*_threshold_ = 0, **Supplemental Figure S5**).

### Hidden Markov models identify three movement states of SPZs in skin

To further quantify the complex motility patterns of malaria SPZs in dermis, we applied hidden Markov model (**HMM**)-based framework, implemented in moveHMM package in R, to cleaned datasets of *P. falciparum* and *P. yoelii* SPZ trajectories from multiple post-inoculation time points (see Materials and methods for detail). Initial application of the method to all trajectories (i.e., assuming that all trajectories are motile, *S*_threshold_ = 0) failed – the model fits never converged. We therefore sub-selected the trajectories with *S* > *S*_threshold_ = 5 *µ*m for further analysis. This selection process removed between 60–86% of tracks, leaving subsets ranging from 52 to 192 trajectories for *P. falciparum* and 100 to 492 trajectories for *P. yoelii* depending on the time point (**Supplemental Table S1** and **Table 4**). We fit different HMMs assuming 1, 2, or 3 states to investigate how many motility states would be needed to accurately describe the trajectory data. Across both species and all time points, the 3-state model consistently outperformed the 2-state model, with ΔAIC values ranging from ~ 800 to nearly 3000 (**Table 4**). This result strongly indicates that SPZ motility cannot be sufficiently captured by a binary classification of “slow” versus “fast” or “circling” vs. “directly gliding” but instead requires at least three latent states to account for the observed heterogeneity (**Figure 8**).

**Table 4:** SPZ trajectory are are best explained by a program with at least three movement states. We show the number of trajectories retained after filtering with *S*_threshold_ = 5 *µ*m (chosen/total, with percentage of removal), AIC values for 2- and 3-state HMMs, and ΔAIC between the models.

| Species | Time post inoculation (mins) | Chosen subset of the experiment data | | AIC (2 state model) | AIC (3 state model) | ΔAIC |
| --- | --- | --- | --- | --- | --- | --- |
|  |  | Chosen tracks/Total | % Removal |  |  |  |
| *P. falciparum* | 5 | 192/1191 | 83.88 | 73969.89 | 71010.67 | 2959.22 |
|  | 10 | 101/255 | 60.00 | 41515.10 | 40169.24 | 1345.86 |
|  | 20 | 115/508 | 77.00 | 49399.30 | 47551.70 | 1847.60 |
|  | 30 | 79/373 | 78.00 | 45502.60 | 44688.80 | 813.80 |
|  | 60 | 52/381 | 86.00 | 27031.50 | 26125.15 | 906.35 |
| *P. yoelii* | 5 | 492/1396 | 65.00 | 86190.09 | 83581.70 | 2608.39 |
|  | 10 | 105/405 | 74.10 | 25860.28 | 24744.53 | 1115.75 |
|  | 20 | 100/560 | 82.10 | 35560.13 | 33426.41 | 2133.72 |
|  | 30 | 149/419 | 59.70 | 35316.83 | 33101.48 | 2215.35 |
|  | 60 | 190/419 | 61.60 | 41988.69 | 40936.60 | 1052.09 |

**Figure 8:**
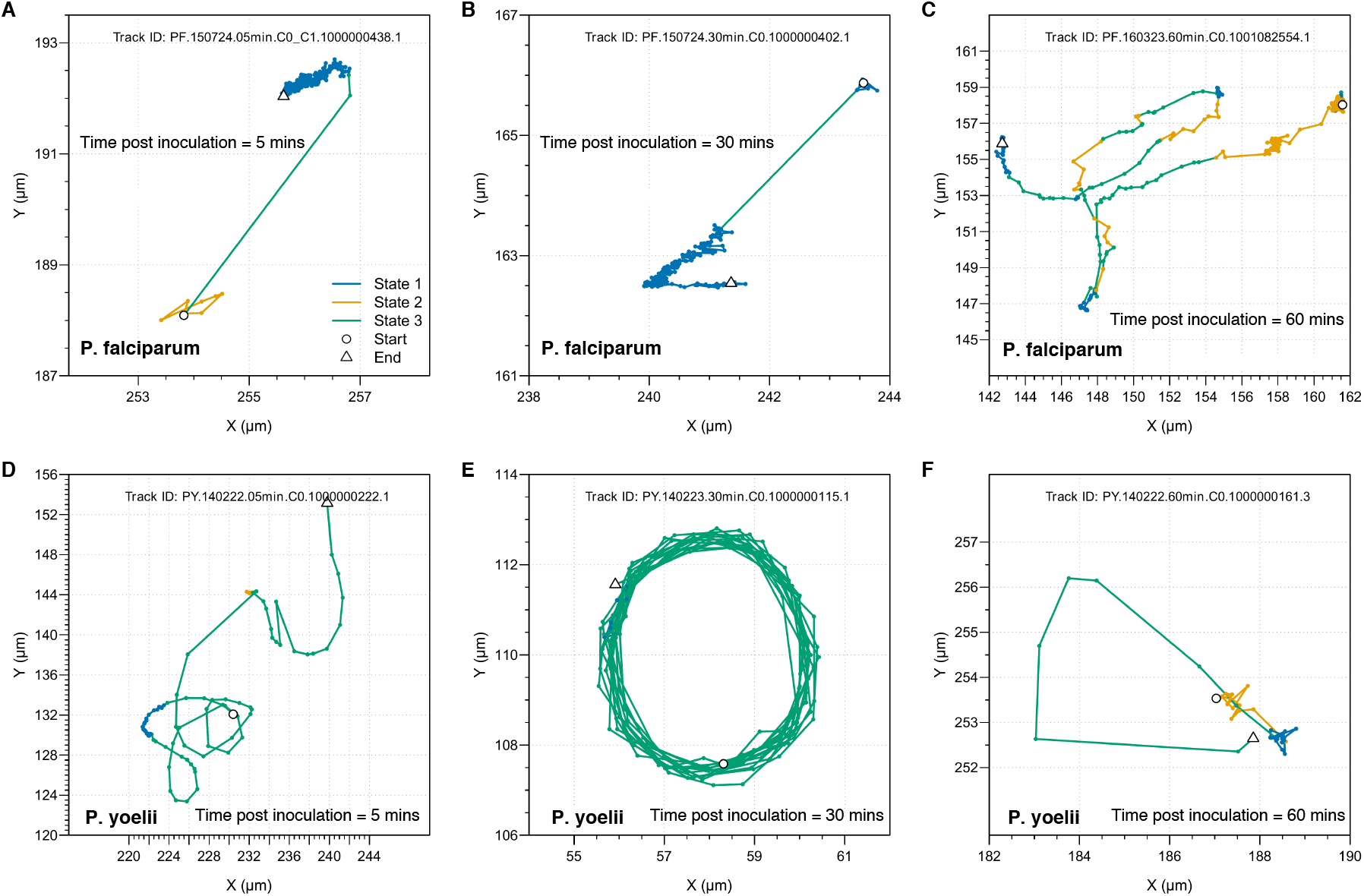
SPZ trajectory data are best characterized by at least three movement modes. We used the R package `moveHMM` to estimate the number of different movement modes that SPZs display in the skin. We used *S*_threshold_ = 5 *µ*m to select for motile SPZs. We show examples of movement trajectories of selected SPZs of *P. falciparum* (**A–C**) and *P. yoelii* (**D–F**), at 5 (A&D), 30 (B&E), and 60 (C&F) min after inoculation. Different colors denote movement modes/states (state 1 — reverse motion with very short step length, state 2 — reverse motion with medium step length, state 3 — gliding/forward movement). Two markers denote the start and end of each trajectory. A model assuming two movement modes fit the data with worse quality based on AIC (**Table 4**). Parameter estimates characterizing the three movement states are given in **Table 5**.

The parameter estimates from the 3-state models (**Table 5**) delineated the following movement modes:

**Table 5:** Parameters of the best fit HMMs with three movement states. We fit the HMM model based on 1, 2, or 3 states to SPZ trajectory data with spatial spread larger than the threshold *S*_threshold_ = 5 *µ*m (see **eqn. (1)**). We list the best fit parameters (the percentage of total travel, step length (mean ± SD, *µ*m), and turning angle (mean and concentration of vM distribution, ^*◦*^)) of the HMM with three states: State 1 represents localized motion with short steps and frequent reversals, State 2 corresponds to reversal movement with intermediate step lengths, and State 3 reflects long, persistent travel with the greatest step lengths and angular concentration. The parameters are for Pf or Py SPZs for different times since inoculation.

| Species | Time post inoculation (mins) | State 1 |  |  |  |  | State 2 |  |  |  |  | State 3 |  |  |  |  |
| --- | --- | --- | --- | --- | --- | --- | --- | --- | --- | --- | --- | --- | --- | --- | --- | --- |
| | | % of travel share | Step length ($\mu\text{m}$) | | Turning angle ($^\circ$) | | % of travel share | Step length ($\mu\text{m}$) | | Turning angle ($^\circ$) | | % of travel share | Step length ($\mu\text{m}$) | | Turning angle ($^\circ$) | |
|  |  |  | Mean | Std | Mean | Conc. |  | Mean | Std | Mean | Conc. |  | Mean | Std | Mean | Conc. |
| *P. falciparum* | 5 | 16.1 | 0.17 | 0.11 | -178.45 | 0.68 | 50.80 | 0.36 | 0.23 | 179.11 | 0.57 | 33.10 | 1.09 | 0.91 | -3.29 | 0.57 |
|  | 10 | 49.90 | 0.29 | 0.20 | 174.58 | 0.46 | 19.90 | 0.71 | 0.37 | 179.01 | 9.96 | 30.20 | 0.87 | 0.68 | -2.44 | 1.41 |
|  | 20 | 17.60 | 0.18 | 0.13 | -179.14 | 0.69 | 55.90 | 0.40 | 0.27 | 179.63 | 0.68 | 26.50 | 1.07 | 0.97 | -4.38 | 0.67 |
|  | 30 | 11.80 | 0.22 | 0.15 | 179.70 | 0.70 | 49.30 | 0.73 | 0.48 | -179.63 | 7.77 | 38.90 | 0.61 | 0.50 | -12.11 | 0.35 |
|  | 60 | 9.80 | 0.16 | 0.12 | 174.12 | 0.46 | 75.90 | 0.49 | 0.34 | -178.77 | 0.70 | 14.30 | 0.68 | 0.58 | 0.22 | 4.08 |
| *P. yoelii* | 5 | 11.30 | 0.29 | 0.21 | -176.75 | 0.34 | 22.80 | 1.03 | 0.79 | -119.77 | 0.08 | 65.90 | 2.22 | 1.92 | 0.91 | 2.76 |
|  | 10 | 12.60 | 0.24 | 0.17 | -169.42 | 0.46 | 35.80 | 0.79 | 0.62 | 170.32 | 0.28 | 51.60 | 1.50 | 1.12 | -4.20 | 3.56 |
|  | 20 | 9.40 | 0.16 | 0.12 | -174.67 | 0.45 | 35.00 | 0.71 | 0.61 | 178.91 | 0.34 | 55.60 | 1.19 | 0.94 | -3.58 | 4.08 |
|  | 30 | 7.30 | 0.16 | 0.11 | -170.32 | 0.41 | 22.30 | 0.63 | 0.53 | 179.55 | 0.32 | 70.40 | 1.37 | 1.11 | -6.40 | 3.95 |
|  | 60 | 6.80 | 0.19 | 0.13 | -173.27 | 0.37 | 29.70 | 0.50 | 0.36 | -178.18 | 0.09 | 63.50 | 1.67 | 1.15 | -3.80 | 2.41 |

- **State 1**: very short step lengths (means 0.16–0.29 *µ*m per sec) and TAs tightly clustered around ± *π*(180^*◦*^), producing frequent reversals and localized displacement. This mode reflects highly restricted motion where parasites remain confined to small dermal regions.
- **State 2**: intermediate step lengths (0.36–1.03 *µ*m means) and ± *π*(180^*◦*^) centric turning angle concentration, indicating meandering or exploratory displacement. Parasites in this state move with moderate persistence, scanning the local tissue environment while still achieving net displacement.
- **State 3**: longest step lengths (0.9–2.2 *µ*m means) and strongly persistent TAs centered around 0, corresponding to directed, long-range travel. This mode captures episodes in which SPZs move most efficiently through the tissue.

Interestingly, in vivo moving SPZs dynamically switch among the three states during a single track (e.g., **Figure 8**). Even within a short trajectory, parasites transition repeatedly among localized reversal, meandering, and persistent travel modes, underlining the stochastic yet structured nature of their motility. Simulations of SPZ trajectories using parameters for 3-state HMM also revealed complex movement patterns and switching between different states (**Supplemental Figure S6**). Taken together, these analyses demonstrate that SPZ movement in skin is best explained as a composite of at least three recurrent motility states but none of these states correspond to a previously defined circling gliding^38–40^. This decomposition captures the diversity of parasite behaviors more faithfully than simpler one or two-state models, provides a quantitative framework for comparing species and temporal dynamics, and highlights the balance SPZs strike between local exploration and efficient forward progression during the critical dermal phase of infection.

## Discussion

In this paper we proposed a novel methodology to classify SPZs as motile or immotile based on the maximal spatial spread that each parasite exhibits during imaging (**Figure 2**). In some cases, such as for Py SPZs it may be possible to define a threshold value *S*_threshold_ that would more rigorously define motile SPZs (**Figure 2**C). We also processed 22 movies on movement of Pf and Py SPZs in murine skins at different times after infection (**Figure 1**, see also deposited datasets on github) and showed that multiple movement characteristics such as final displacement, mean speeds, mean squared displacement and turning angles strong depend on the chosen *S*_threshold_ (**Figures 3–7** and **Tables 2 and 3**). In particular, higher values of *S*_threshold_ made change in average speed of Pf or Py SPZs less dependent on the time since inoculation. Inference of the movement patterns, based on early MSD slopes or fits of mixture of vM distribution to TA data, was also dependent on which tracks are deemed motile; defining motile SPZs as those with *S* > 5 *µ*m changed classification of Py SPZs from largely reverse movements to primarily forward movements (**Figure 7**B&D). We also showed how using a hidden Markov model-based analysis of SPZ trajectories allows to rigorously determine different movement modes/states, and for SPZs in skin, we found 3 different movement states (**Figure 8** and **Tables 4 and 5**). Our analysis also revealed differences in movement of Pf and Py SPZs in murine skin with Pf SPZs clearly showing slower speeds and less persistent displacement (**Figures 4 and 5**).

Our results on quantifying movement patterns of Pf and Py SPZs when using a relatively conservative MSS threshold (*S*_threshold_ = 5 *µ*m) are broadly consistent with some of the previous findings but are not with some others. In particular, we found small decline in final displacement or average speed of Py SPZs with time since inoculation as previously ^17^; however, in contrast with Hopp *et al*. ^17^ study we found much slower decline in average speed of Pf SPZs with time since inoculation (**Figure 4**C, our slope *m* = − 3.3 × 10^*−*4^*/*min and *p* = 0.6 vs. previously found *m* = − 20.0 × 10^*−*4^*/*min with no p value reported). We also found that Py SPZs exhibit consistently higher average (or instantaneous) speeds as compared to Pf SPZs for the same time since inoculation (*p* « 0.01 for most comparisons); that is inconsistent with previous observation that found no significant difference between average speeds of Pf vs. Py SPZs for most times since SPZ inoculation^17^. Overall change in MSD with time suggests that both Pf and Py SPZs undergo correlated random walks, a subset of Brownian walks, and this is different from the previous work that analyzed step length distribution and suggested that SPZ do not undergo Brownian walks^13^. Finally, there have been multiple studies classifying SPZ movement in vitro and in vivo^8,38–42^. While there have been differences in what types of movement modes different studies identified, “circling gliding” or simply circling ^38^ has been a consistent mode of SPZ movement. Our application of a new methodology to detect hidden movement states suggested that the data are best explained with at least 3 states but none of them directly corresponded to circling (**Figure 8**). Because many other studies used very distinct criteria to define motile tracks (**Table 1**), it is harder to compare our results with other previous observations. Our results therefore suggest that some variation across published reports may arise not only from biological differences in parasite species, host environment, or imaging system, but also from differences in sub-selection of tracks deemed as motile.

Our analysis of SPZ track suggests that both SPZ species undergo correlated random walks that allow faster than Brownian short-term displacement but also space exploration due to turning and circling. The goal for skin-deposited SPZs is to find and invade blood vessels but at present there is limited evidence that SPZs are attracted to the vessels and seemed to find them by random/unbiased movement^8^. This is in sharp contrast to neutrophils, white blood cells, that tend of display strong bias in movement towards a site of injury or location of a parasite in the skin^43,44^. Liver- or brain-localized CD8 T cells also undergo correlated random walks ^22,23^, and only a small proportion of CD8 T cells, specific to Plasmodium antigen, display minor attraction to the SPZ in the liver^30^. Thus, despite different mechanisms regulating motility of SPZ and immune cells, some characteristics of their movement (e.g., correlated nature of walks) are quite similar. HMM-based approaches provide an additional framework for identifying state-dependent movement programs across different biological systems. Thus, some motility metrics can reveal common movement features, while differences in speed, turning angle distribution, MSD slope, and movement-state composition reflect distinct mechanisms of motility and the environmental constraints experienced by each cell or pathogen type. This broader applicability makes our analytical framework relevant beyond Plasmodium SPZ to a wide range of motile biological systems.

Several limitations should be considered when interpreting the results of this study. First, although we had access to the total of 97 movies, we only processed 22 of these, in part, due to large amount of work required to carefully trace positions of many SPZs in each movie. While this subset contained sufficient trajectories to evaluate the effects of motility-threshold selection, our selection of the movies to analyze may not be fully random, and this could in part explain differences with previous results^17^. Second, parasite trajectories were generated using automated tracking in Imaris and subsequently evaluated through manual inspection to ensure correspondence with actual parasites. Manual correction may introduce biases in defining which trajectory corresponds to which SPZ especially when there are hundreds of parasites and they move in and out of the imaging volume. Semi-automated approaches have been used in previous sporozoite tracking studies but their validity remains to be explored further ^17,18^. Third, the trajectory datasets contain occasional gaps in time frames, which commonly occur in time-lapse microscopy due to temporary loss of signal or tracking discontinuities^45,46^. We have performed our analysis with cleaned trajectory data but this typically results in more tracks with fewer time frames per track. Fourth, sporozoite migration occurs in three dimensions, whereas trajectories analyzed here are derived from two-dimensional Z-projections. Consequently, measured displacement and velocity may underestimate true parasite movement^4,13,47^. Finally, the datasets were generated using syringe inoculation rather than mosquito bite, which may influence parasite dispersal and early migration patterns in the dermis^4,7,27^.

Our work opens avenues for future research. There is a need for a broader discussion of what constitutes a motile SPZ in vivo. While we propose to use maximal spatial spread threshold to define a given SPZ as motile, other metrics (e.g., based on speed) may be also appropriate. We found that intermediate thresholds (*S*_threshold_ = 5 – 7 *µ*m) yield stable estimates of speed and MSD slopes while preserving adequate sample size—representing a practical compromise between inclusivity and specificity. Whether similar threshold values would work in other studies would be important to investigate. Incorporating fully three-dimensional imaging approaches may further improve the accuracy of motility measurements and provide a more complete description of parasite migration dynamics within the dermis^4,13,27^. In addition, integrating spatial information on dermal blood vessel locations could add important biological context to trajectory analyses, allowing parasite movement patterns to be interpreted relative to potential vascular entry points^8,17^. Such spatially informed analyses may help distinguish exploratory migration from directed movement toward blood vessels and provide deeper insight into how sporozoites locate and access the host vasculature. Together, these approaches may facilitate the development of more rigorously defined metrics for characterizing sporozoite motility and enable more consistent comparisons across studies.

## Abbreviations

MSS: maximal spatial spread
Pf: Plasmodium falciparum
Py: Plasmodium yoelii
SPZ: sporozoite
MSD: mean squared displacement
TA: turning angle
vM: von Mises
HMM: hidden Markov model
LRT: likelihood ratio test
AIC: Akaike Information Criterion.

## Data sources

The imaging data has been provided by Dr. Christina Hopp from their previous publications^13,17^; the master spreadsheet containing sporozoite coordinates is available on github: https://github.com/drsbiswas/Plasmodium-SPZ-Movement-in-skin/releases/tag/Source-code-and-dataset

## Code sources

All analyses have been performed primarily in Rstudio (ver 4.3.1) or Mathematica 12.2.

## Ethics statement

No new experiments have been performed.

## Author contributions

E.H. and V.V.G. conceived and designed the study. E.H. analyzed the imaging data and generated SPZ trajectory dataset. E.H. also developed the computational analysis pipeline and conducted initial statistical analyses, including mean squared displacement and turning angle distributions, and documented the workflow. V.V.G. performed von Mises distribution fitting for turning angle data. S.B. performed hidden Markov model inference. S.B. and V.V.G. interpreted the results and developed the conceptual framework of selecting motile tracks using maximal spatial spread. S.B. conducted statistical analyses across varying spatial spread thresholds and wrote the initial draft of the manuscript. All authors contributed to editing and revising the manuscript and approved the final version. V.V.G. supervised the study.

## Acknowledgments

We would like to thank Christina Hopp for sharing movies from their published studies, and Photini Sinnis and Tim Anderson for comments on earlier versions of the paper. We would like to thank Anderson King, Kailynn Deck, and Whitney Griffin, undergraduate students at the University of Tennessee, Knoxville, who helped with initial analysis of the imaging data with Imaris, Nikki de Vries from Texas Biomed for making supplemental videos for the paper. This work was supported in part by the NIH grants R01GM118553 and R01AI158963 to VVG.

## Supplemental Information

**Movie 1:**
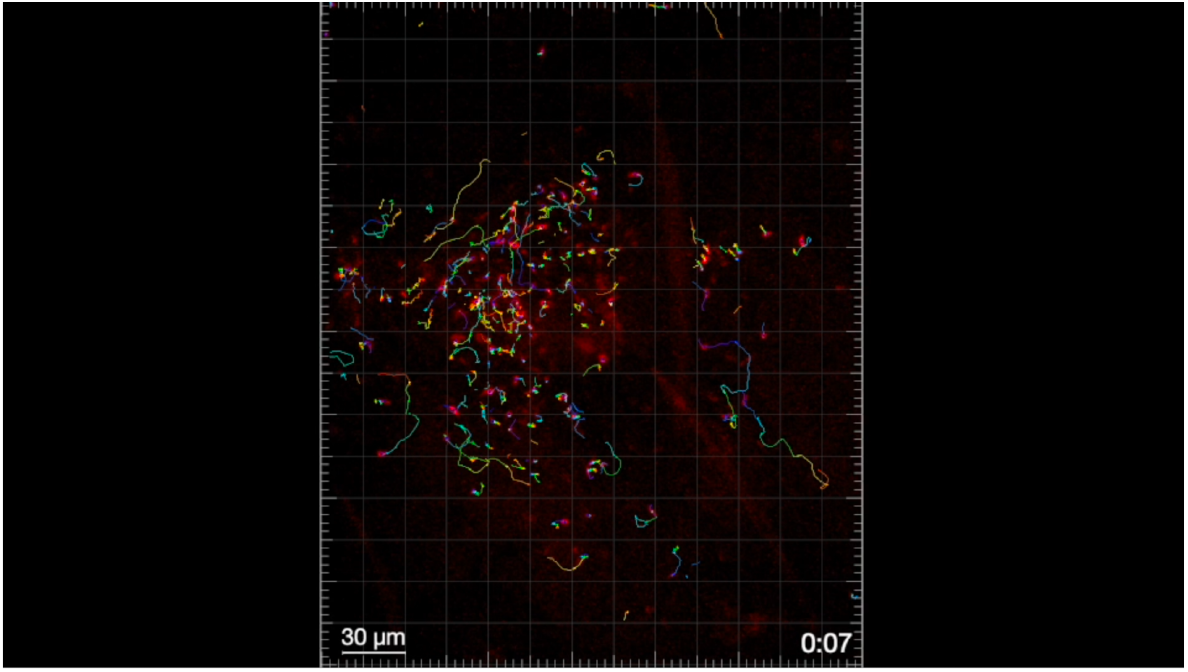
Quantifying trajectories of Pf sporozoites in skin. We used Imaris to rigorously quantify positions of Pf sporozoites (see Materials and methods for detail). Bar scale is 30 *µ*m. The volume of 512 × 512 × 2 *µ*m was scanned every 1 sec. Time is shown in min:sec.

**Movie 2:**
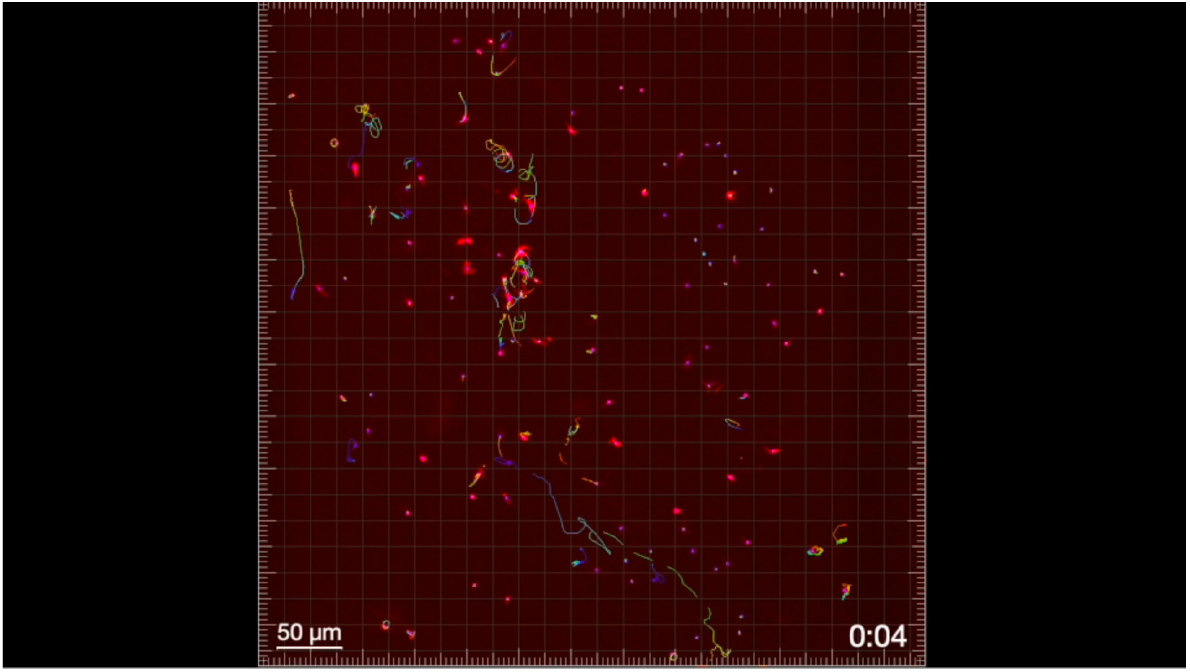
Quantifying trajectories of Py sporozoites in skin. Same as **Movie 1** but for Py sporozoites with scale bar being 50 *µ*m.

**Supplemental Figure S1:**
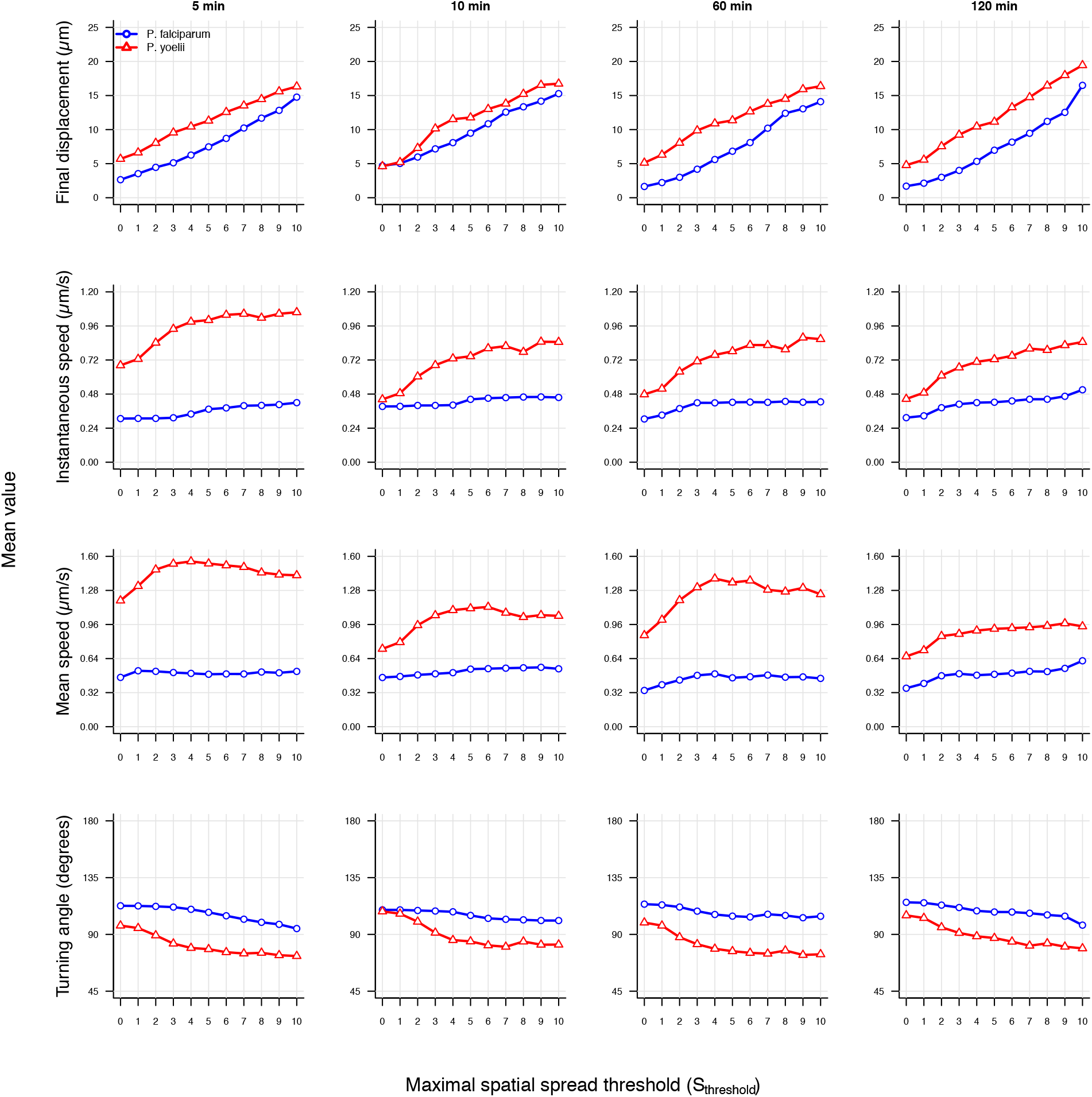
Increasing the maximal spatial spread threshold yields higher mean displacement and speed values for *P. falciparum* and *P. yoelii* SPZs. We show the average final displacement, instantaneous speed, and mean speed, for each displacement cutoff (*S*_threshold_ = 0–10 *µ*m) and different times after SPZ inoculation into the skin (5, 10, 60, and 120 min). Metrics were derived from trajectories retained after spatial spread filtering and time-step cleaning of Imaris-processed datasets.

**Supplemental Figure S2:**
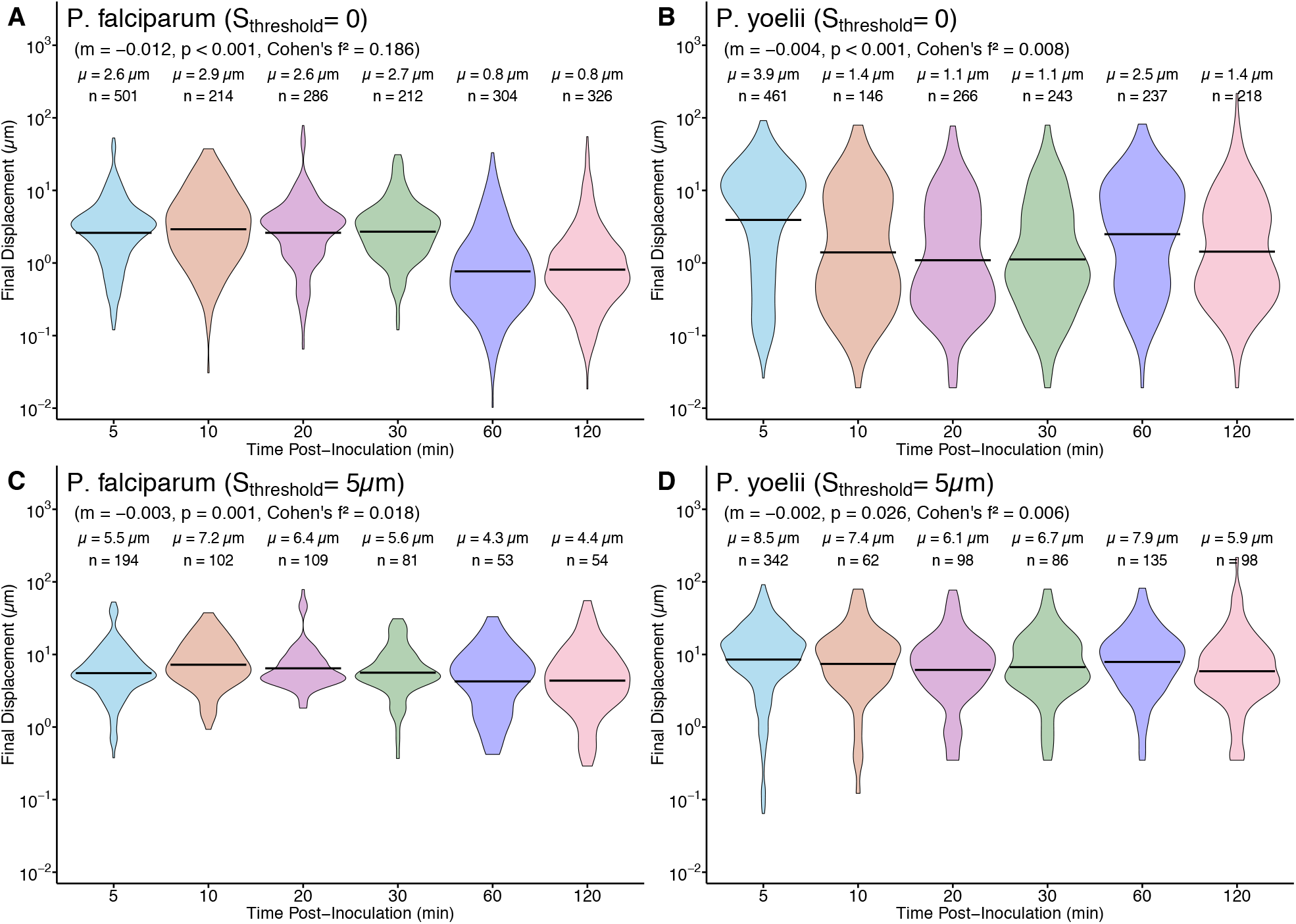
Final displacement of Plasmodium sporozoites is larger for unclean data and only weakly depends on the time since inoculation. We performed similar analysis as in **Figure 3** but for original/unclean data that includes missing time frames for some SPZs (see Materials and methods for detail of data cleaning). Final displacement was different between Pf and Py at 5, 60, and 120 minutes post-inoculation (5 min: *p* < 0.001, Cohen’s *d* = 0.651; 60 min: *p* < 0.001, Cohen’s *d* = 0.707; 120 min: *p* < 0.001, Cohen’s *d* = 0.336).

**Supplemental Figure S3:**
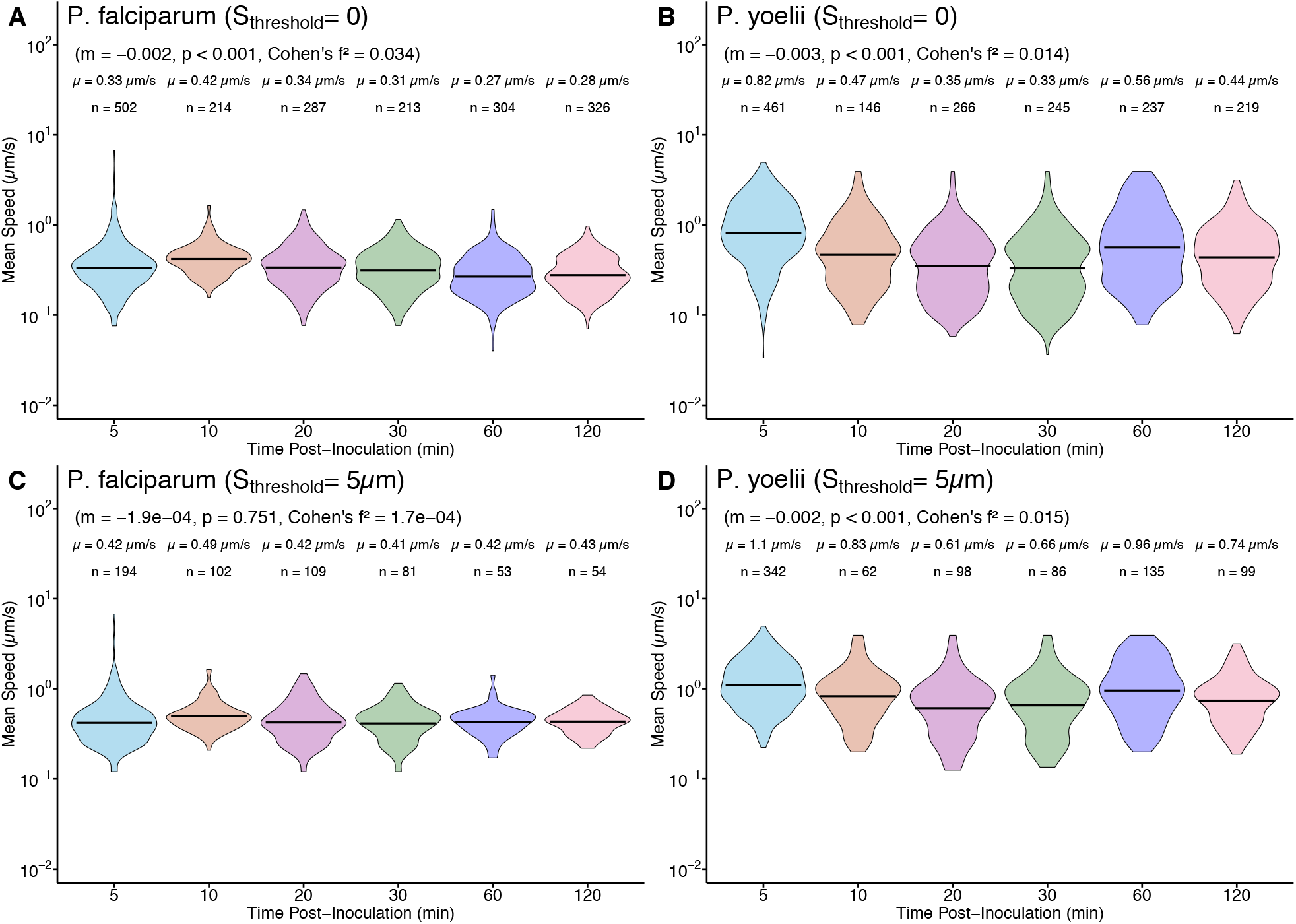
Mean speed of Plasmodium sporozoites weakly depends on the time since inoculation in original/unclean data. We performed similar analysis as in **Figure 4** but for original/unclean data. Mean speeds were different between the two species at all times post inoculation (5 min: *p* < 0.001, Cohen’s *d* = 1.074; 10 min: *p* < 0.001, Cohen’s *d* = 0.477; 20 min: *p* < 0.001, Cohen’s *d* = 0.280; 30 min: *p* < 0.001, Cohen’s *d* = 0.340; 60 min: *p* < 0.001, Cohen’s *d* = 1.00; 120 min: *p* < 0.001, Cohen’s *d* = 0.840).

**Supplemental Figure S4:**
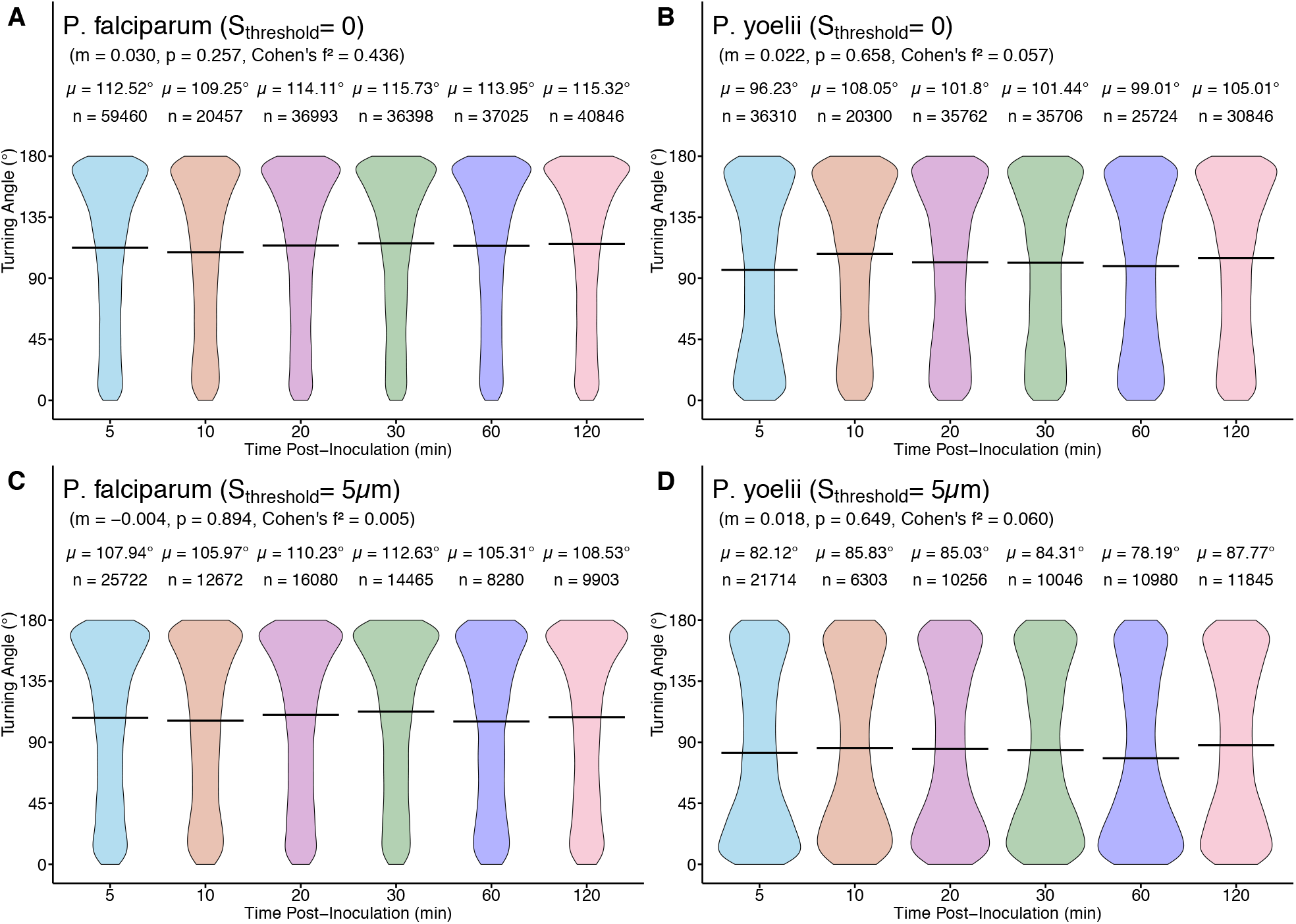
Similar proportions of clock and counterclock turns of Plasmodium SPZs are independent of time since inoculation. For each SPZ trajectory in the cleaned dataset for Pf (**A**) or Py (**B**) we calculated TAs by taking into account whether turning occurs clock- or counter-clock wise. We show the data as violin plots with the number of angles (*n*) and average value of the mean speed for all SPZs (*µ*).

**Supplemental Figure S5:**
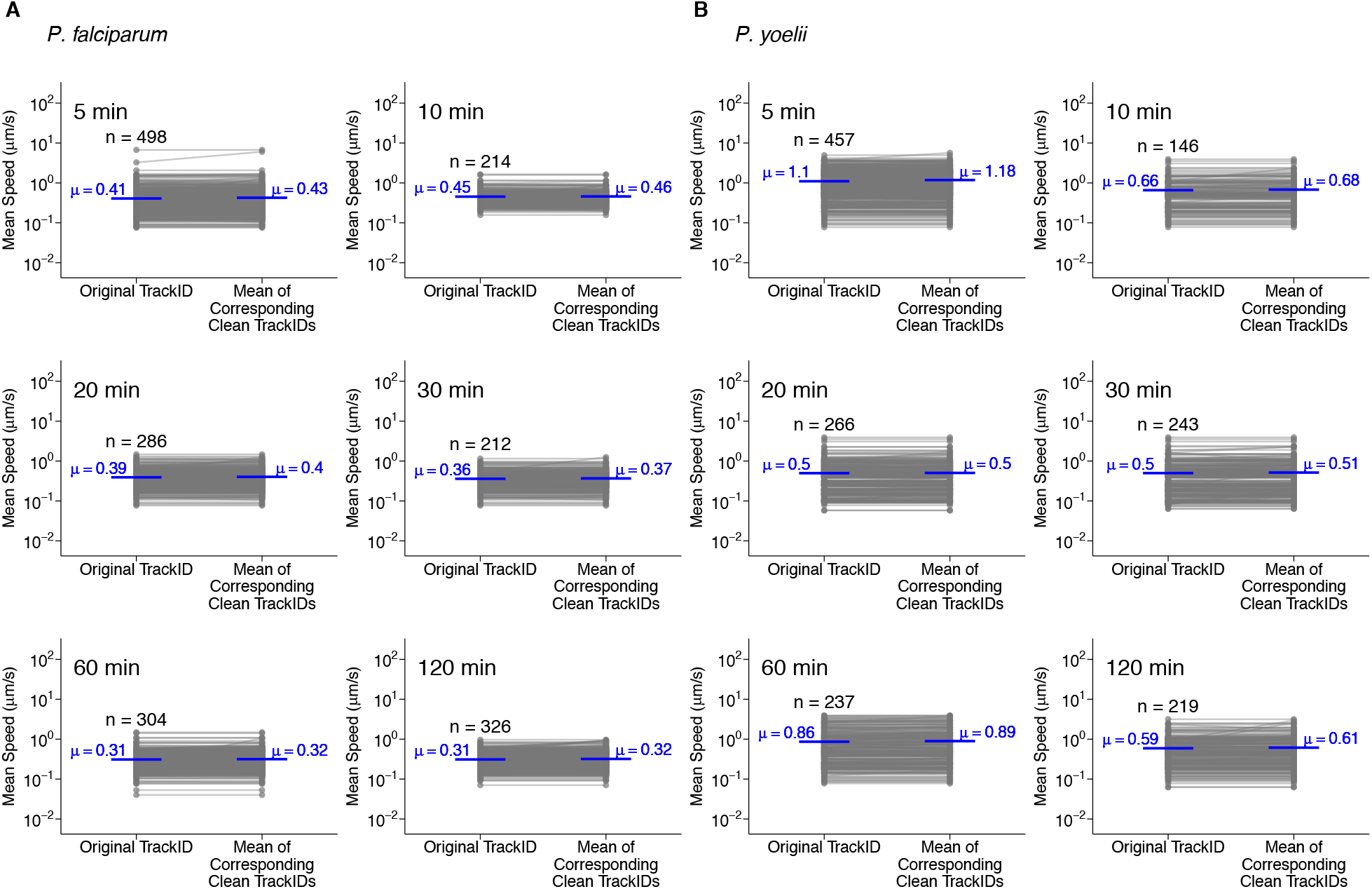
Cleaning of trajectory data has a minimal impact on mean speed of Plasmodium sporozoites. For every SPZ trajectory in the original data for Pf (**A**) or Py (**B**) we generated a set of clean trajectories for which every SPZ position was continuous in time (see Materials and methods for detail of how cleaned trajectory data were generated). We then calculated the mean speed for every SPZ trajectory in the original dataset and mean speed of all split tracks corresponding to the same original track ID. Specific times after SPZ inoculation at which speeds were measured are indicated on individual panels. We found higher speeds for cleaned data for Pf at 5 min (*p* = 0.006, Cohen’s *d* = 0.038), 20 min (*p* = 0.004, Cohen’s *d* = 0.048), 60 min (*p* = 0.047, Cohen’s *d* = 0.032), and 120 min (*p* < 0.001, Cohen’s *d* = 0.057). For Py, we found higher speeds for cleaned data at 5 min (*p* < 0.001, Cohen’s *d* = 0.092), 10 min (*p* = 0.024, Cohen’s *d* = 0.034), 30 min (*p* = 0.011, Cohen’s *d* = 0.029), and 60 min (*p* = 0.001, Cohen’s *d* = 0.034).

**Supplemental Figure S6:**
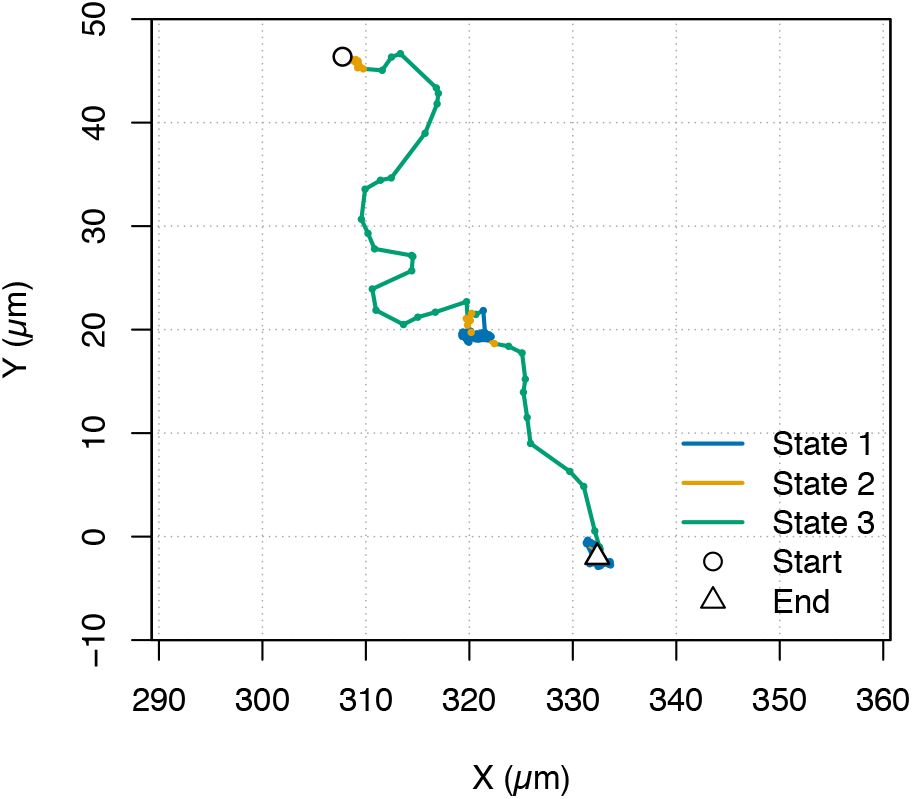
Simulated trajectory reproduces key features, both the spatial heterogeneity and state-switching dynamics of observed sporozoite motility. By using parameters estimated using 3-state HMM (see **Figure 8** and **Table 5** for Py SPZs at 5 min post inoculation) we simulated a SPZ movement using the `simData` function in `moveHMM`. The number of steps were chosen randomly from the all the trajectories in dataset and is anchored at a random starting position. Segments are colored by inferred states.

**Supplemental Table S1:**
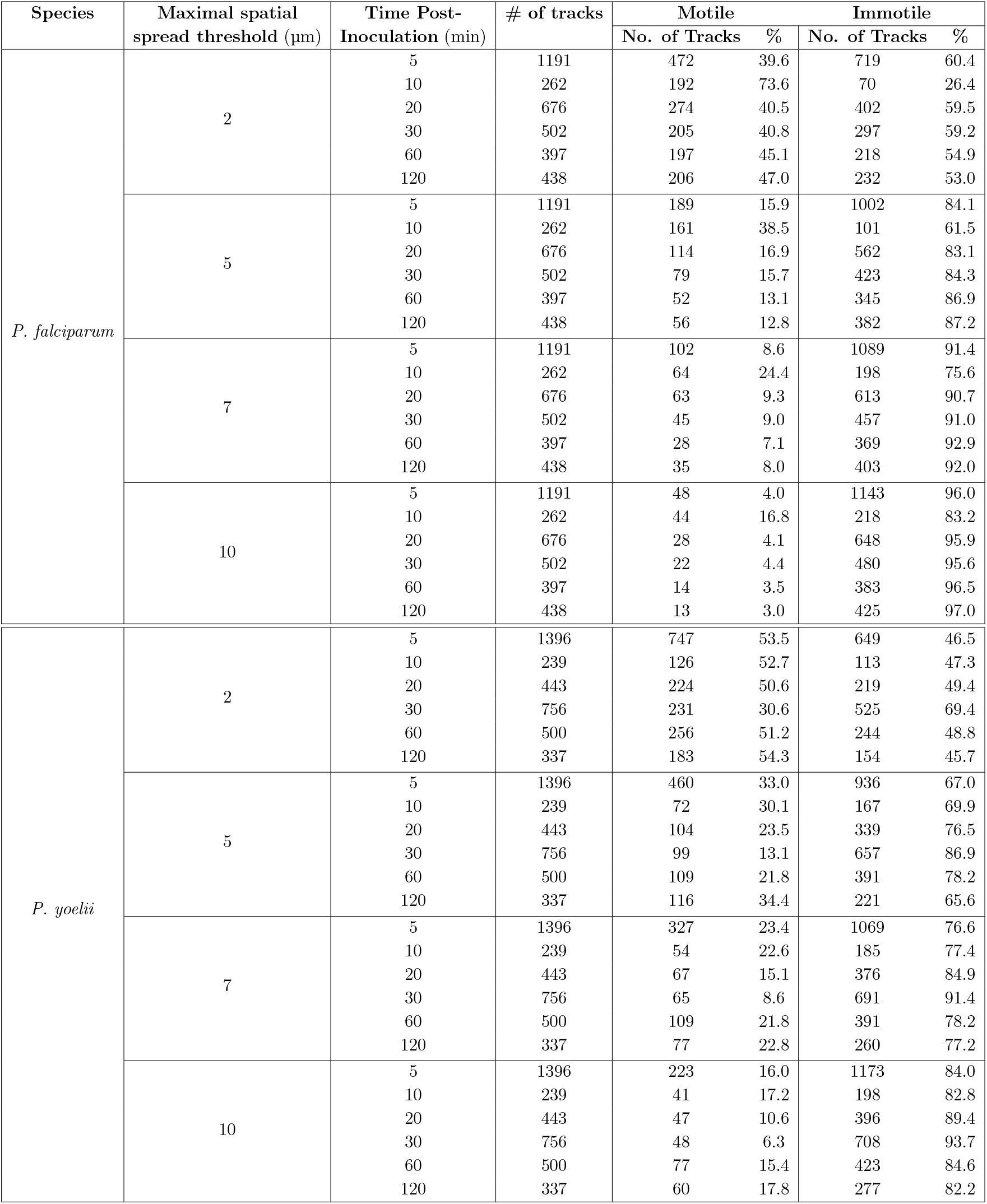
Larger spatial spread thresholds reduce the number of trajectories defined at motile in both *P. falciparum* and *P. yoelii*. We show the total number of trajectories and the counts and percentages of motile and immotile tracks obtained for each spatial spread threshold value (*S*_threshold_ = 2–10 *µ*m) and time point after inoculation (5–120 min). We cleaned the trajectories (by splitting trajectories that had missing time frames) and filtered using the spatial spread threshold (see **eqn. (1)** and Materials and methods for detail).

